# The balance stabilising benefit of social touch: influence of an individual’s age and the partner’s relative body characteristics

**DOI:** 10.1101/2024.11.21.624637

**Authors:** Katrin H. Schulleri, Dongheui Lee, Leif Johannsen

**Author notes:** **Corresponding author:** Leif Johannsen, **Email:**, Address: Cognitive and Experimental Psychology, Institute of Psychology, RWTH Aachen University, GermanyJägerstrasse 17/19, D-52066 Aachen, Germany.

## Abstract

Interpersonal touch (IPT) is a successful strategy to support the stability of and individual’s body balance during a multitude of activities in daily life, including physical education and therapy. Despite common practice, however, the influence of an individual’s anthropometry and other personal characteristics, such as age, balancing skills, motor experience, and sex as well as interindividual differences in these characteristics between interaction partners on the balance stabilising benefit of social touch is unknown.

We assessed an individual’s balance stability and change due to IPT provision during single-legged stance in 72 pairs (age range 4 to 63 years) under four sensory conditions: with or without vision in combination with IPT or without. Two participant subgroups were created: one of more vulnerable with low stability and one of more stable, mature participants. Best fitting multiple linear regression models, including moderating variables, for explaining the benefit of IPT in each visual condition indicated that without vision, an individual’s benefit of IPT was determined by their balancing skill and the partner-related difference in balancing skill but not by any other factors or partner-related differences. Especially vulnerable individuals improved considerably with IPT when vision was unavailable. With vision complementing IPT, however, an individual’s age-dependent motor developmental potential became an additional moderating factor.

These findings indicate that the extent to which IPT is benefitting mutual balance stabilisation does not depend on biomechanical factors. Instead, the IPT benefit emerges as a product of both partners’ sensorimotor capabilities moderated by a person’s motor developmental potential when visual feedback could be utilised. We discuss a theoretical framework that accounts for the observed dependencies of the effect of haptic social support on balance control.

**Significance Statement:** This work contributes to a better understanding of the factors influencing the benefit of interpersonal touch for balance stabilization across a wide age range from children to senior adults. It is the first work that investigates the role of both individual factors and relative differences between interaction partners on the amount of benefit of IPT and takes moderating factors such as age-related motor developmental potential, sex, and body mass index into account. Future studies can build on this work when selecting or simulating an optimal partner, especially for individuals with postural instability, for example due to age or sensorimotor deficits.

## Introduction

Falls are a leading cause of injury-related deaths worldwide. One of the main causes of falls is impaired body balance (1). Several factors are known to influence the stability of body balance, such as age (2–4), sex (5, 6), and anthropometry (7–9). Balance control can be facilitated by tactile feedback (10–13). Just a light touch is sufficient to inform about any motion of one’s own body relative to an earth-fixed reference point and thereby improve stability through optimised balance adjustments (14–16). When physical contact is kept with an environmental reference, internal representations, such as the body schema, are involved in the localization of the contact and its relative motion and considering the context of specific body postures and the limbs of the body, localization occurs in an egocentric frame of reference (17).

The stabilising benefit of external light touch on balance stability generalises to social interactions. Interpersonal balance support is frequently observed in daily life, such as when providing support to a frail person in a clinical setting or to a person with reduced balance stability because of challenging or inadequate footwear (e.g. high heels). Interpersonal touch (IPT) with a dynamic reference such as another human also leads to enhanced balance stability and spontaneous interpersonal postural coordination (“sway entrainment”), possibly due to improved state estimation by augmented sensory feedback about own sway dynamics and perception-action coupling with an intention to minimise perceived interaction force fluctuations. Passive exposure to the recorded sway dynamics of another individual via haptic force feedback, however, does not result in sway reductions in the way it is normally observed during contact with an actual human partner (18). Therefore, sway reduction with IPT may reflect a mutually adaptive process between two contacting individuals and not just the reception of additional haptic information (19). Furthermore, the effect of IPT does not seem to be the sole result of a mechanical coupling between both individuals but instead may represent the effect of mutually shared sensory information (20). Thus, the social context during balancing with IPT seems to have an important influence in addition to mechanical factors.

From a developmental perspective, it is a truism that the younger the age of a person and the less motor experience they possess, the more they are in need to acquire fundamental motor capabilities. Nevertheless, they have the greatest potential for extending and optimising their motor repertoire in multiple directions. During early motor development, haptic interactions with another individual play an important role for the development of postural control and the involved body representations (21). Bremner (22) characterised the acquisition of multimodal body representations as an interface between an individual’s body and the external environment. Further, Bremner and Spence (21) regarded these representations and the ability to utilise touch fundamentally for the development of a subjectively experienced self. Early-stage toddlers seem to be quite susceptible to haptic inputs that convey self-motion information gained from environmental contact. For example, Chen and colleagues (23) observed that light haptic contact stabilised balance markedly during the transition period when toddlers began to take their first independent steps. They interpreted this as an indication of progressively refined internal representations of their own sway dynamics during standing and walking (23). Ivanenko et al. (24) demonstrated in toddlers how unilateral single hand support with a parent improves postural stability in terms of reduced trunk sway, sideways hip motion as well as step width. These benefits were seen as an indication of increased toddlers’ stepping confidence but may also be interpreted as parental facilitation of the acquisition of locomotor skills.

In children, the processing of sensory information, internal state estimation, and motor control are affected by greater amounts of internal noise (25–27). Proprioceptive sensory information appears to have the greatest influence on balance stability followed by vision and tactile feedback (28). Multisensory integration and reweighting changes with age and is optimised in more mature young individuals (29–35). Furthermore, internal representations and feed-forward, predictive control are developed gradually (36, 37), while cognitive control continues to contribute to a greater extent than in adults (38). Since children possess immature multisensory integration, less precise body representations, are less stable and more susceptible to haptic information, one could expect that younger individuals benefit more from IPT. In addition, their developmental potential, meaning the length of their future developmental path, may raise the contextual relevance of their individual balancing skill in relation to the balancing skill of an interaction partner.

In old age, the developmental progression towards sensorimotor maturity observed in children and adolescents seems to be reversed. Noise in sensory feedback increases due to age-related deterioration in sensorial acuity (39), and sensory processing may slow down because of demyelination (40). Therefore, older adults may begin to rely more on the integration of redundant information from multiple different sensory channels, such as vision and somatosensation (41). Thus, multisensory integration is altered with older age (42), e.g. visual dependency for balance control increases as the reliability of proprioception and vestibular sensation decreases (43). Finally, as muscle strength weakens with age, potentially less optimal motor control strategies with a lower muscular effort are adopted (39). All in all, one can argue that the motor developmental potential of older adults is minimal compared to children, albeit not zero given enough daily exercise and practice to uphold their level of fundamental motor capabilities.

The developmental changes in childhood and adolescence and the deterioration in older age can be described as a U-shaped relationship (44) between age and balance skills across the range of 5 to 100 years of age, with a valley floor between 20 to 40 years (resembles an L-shaped function from 5 to 40 years) (3, 45, 46). During childhood and adolescence, postural control improves with age and variability in balance performance decreases as stability increases (47) until the age of around 20-25 years, when a plateau is reached. After around the age of 45-55 years, body sway may begin to increase again, and efficiency of postural control deteriorates with older age (4, 46).

A U-shaped relationship has also been observed between body sway and body mass index (BMI) as a derived measure combining anthropometric characteristics such as body height and weight. Lee et al. (8) investigated the relationship between balance stability and BMI in a large cross-sectional study of community-dwelling older adults between 65 and 92 years of age. An excessively increased BMI is associated with balance instability and increases an individual’s risk of falling. Furthermore, individuals with extremely low BMI, for example, due to eating disorders such as anorexia and bulimia, also showed reduced balance stability compared to individuals with normal BMI (48, 49). Interestingly, individuals with extreme BMI seem to demonstrate inadequate multisensory integration (50) and greater sensitivity to external tactile stimulation (51). Consequently, we would expect individuals with extreme BMI to benefit more from IPT. Moreover, we would expect the importance of the own balancing skill and the difference in balancing skills compared to their interaction partner for the benefit of IPT to be moderated by extreme BMI.

In addition to the state of sensorimotor development, ageing, and anthropometry, the gender or sex of an individual seems to have a distinct influence on balance control, too (52). Evidence exists that sensorimotor control and the integration of external feedback may be organised differently in males and females. Specific moderating factors may be weighted differently between the genders. For example, studies on motion sickness (53) and ‘mal de debarquement’ syndrome (54) found females to be more susceptible to indications of imbalance during multisensory conflict. It has been hypothesised that central processes in multisensory integration, such as visual-vestibular interactions, are not just task-dependent but also may differ between sexes (or gender identities; 55, 56). Thus, one could expect sex to influence the benefit of IPT as well and affect the role of an individual’s and partner’s balancing skill on the benefit of IPT.

The influence of individual characteristics, such as age, anthropometry, and sex, on the benefit of IPT is unknown, and it is unclear to what extent also an interaction partner’s personal characteristics play a role regarding the benefit of IPT. Consequently, the question arises if someone like an “optimal” or most suitable IPT partner can be defined from whom one would benefit the most in terms of balance stabilisation? An asymmetry is known where the less stable individual in a pair shows much greater improvements than the more stable partner. The effects of the tactile interpersonal interaction on balance stability could be mediated by biomechanical factors such as the weight, height, and the intrinsic stability of an individual and their partner. For example, a person in bipedal stance may improve balance stability of a person in less stable Tandem Romberg stance by damping body sway.

In the present study, we circumvented these biomechanical confounds by adopting an intrinsically much more challenging standing posture, single-legged stance, in which performance seems to be strongly age-dependent (57) and where even a person with little body mass could easily perturb the balance of a person with a greater amount of inertia.

Thus, we used this postural context to tease apart possible confounding factors determining the benefit of IPT on balance stability. We adopted a broad approach to investigate the potential factors influencing the benefit of IPT. We expected a greater benefit of IPT to be explained by an individual’s balancing skill, by sex, by age (in terms of motor developmental potential), and by anthropometry (BMI). Furthermore, the benefit of IPT was also assumed to depend on the relative differences in balancing skill compared to the partner’s skill (i.e. greater benefit with a more stable partner), as well as in sex, age (greater benefit with a more mature partner) and BMI. Figure 1 summarises our conceptual framework of interactions between parameters.

**Figure 1.**
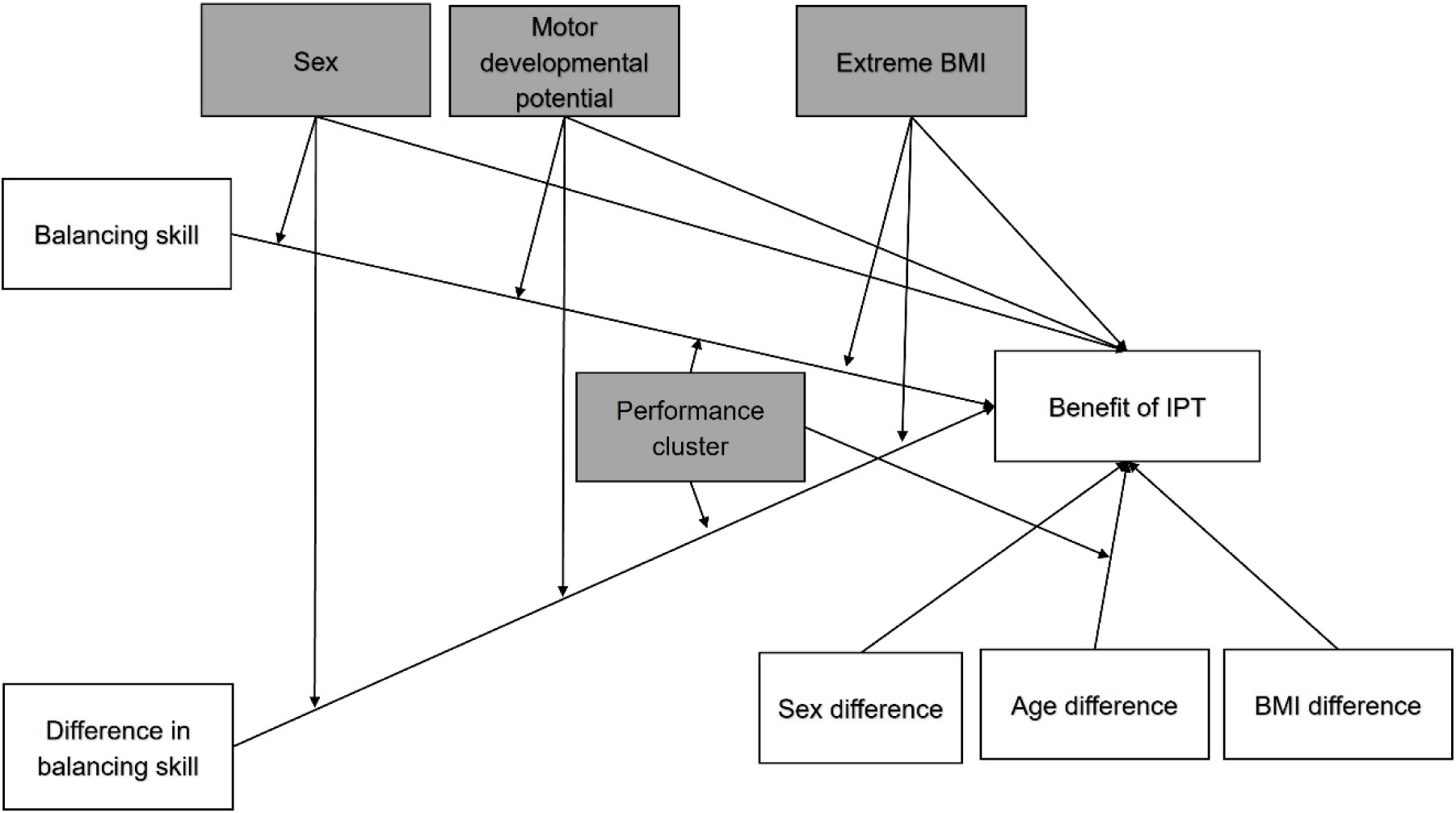
Conceptual model of the influential factors on the benefit of IPT, sex, age-related motor developmental potential (mean-centred age inverse), extreme BMI (mean-centred BMI squared) and performance cluster assignment (vulnerability of individuals) moderate the effects of an individual balancing skill, proportional interindividual differences in balancing skills, and interindividual differences in sex, motor developmental potential, and BMI. IPT: interpersonal touch; BMI: Body mass index.

## Methods

### Participants

Within the period from September 1^st^ to November 30^th^, 2019, one hundred and sixty-two participants were recruited as an opportunity sample at public science festivals showcasing research undertaken at the Technical University of Munich. Individuals approaching our public display received an explanation of human sensorimotor control of body balance and the functioning of a barometric platform for stance and gait analysis and were invited in random pairs to take part in this study. Individuals in pairs could be related to each other (partners, parent and child, friends) or be unacquainted. The investigation was carried out in accordance with the Declaration of Helsinki on Ethical Principles for Medical Research Involving Human Subjects and was approved by the medical ethical committee of the Technical University of Munich (2019-248-S-SR). All participants and their parental guardians, when participants were underaged, gave verbal informed consent. As the data collection was conducted in public the obtaining of consent was witnessed by any onlookers. Participants were excluded from subsequent data analysis, but not from taking part, if they reported pre-diagnosed sensorimotor impairments known to affect control of body balance, such as stroke or polyneuropathy. The experimental procedure was explained to each participant, and they could refuse to participate at any time. Participants remained anonymous throughout data collection and did not provide any personalised information that would make them identifiable. Only a participant’s age, gender, body height and weight were recorded. Participants did not receive any evaluation or feedback about their balancing performance.

### Experimental design

The control of body balance can be challenged by a small base of support, e.g. as in a Tandem-Romberg stance or in single legged stance. Standing on one leg is a good indicator for an increased risk of falling (58) and is dependent on balance-specific feedforward and feedback sensorimotor control skills that are observed in experienced dancers (59). Therefore, we adopted a single-legged stance paradigm in the present study. Participants were instructed to stand side-by-side and orthogonal to the length of the pressure plate in a single-legged stance with stockinged feet on their preferred leg. Due to the relative difficulty and unstable nature of a single-legged stance (compared to a normal bipedal stance), we expected that also a taller and heavier individual’s body sway could be destabilised severely by only slight perturbations imposed by an interaction partner. Therefore, we assumed that the body sway of shorter and lighter individuals would not be stabilised during IPT due to mechanical damping by a taller and heavier partner.

Data acquisition took place in public, but participants had their backs turned towards any potential onlookers by facing a blank, white wall at a 2-metre distance. A pressure plate two metres wide (Zebris FDM 2; single sensor dimensions: 8.46 mm * 8.75 mm; 240 * 64 sensors) was used to record the foot pressure distributions of a pair at a sample rate of 60 Hz. Four single trials of 20 seconds duration, one for each stance condition (Eyes open and Eyes closed both with and without IPT), were acquired so that the entire procedure lasted 5 minutes for each pair. The four conditions were tested in random order. For the administration of the interpersonal touch, participants in a pair were instructed not to grasp each other’s hands but to rest their fingertips against each other, as reported by Johannsen et al. (13). Participants were also asked to keep the same arm posture in all four trials, and the hand of the extended arm was usually held in supination. The demands of single-legged standing, however, could result in participants deviating from the target posture, for example, when trying to rebalance themselves during a phase of instability. When interpersonal touch was available, the individual on the left held the right hand in pronation to contact the fingers of the person on the right from above. Participants were instructed to remain relaxed without speaking during each trial.

### Data reduction

During post-processing, the pressure plate matrices for each data frame were divided into one area for each participant’s footprint. From each trial, the longest period of static standing was visually determined and manually segmented. In the best of cases, the longest period of static standing covered the entire length of a trial. In those trials in which one of the two participants lowered their initially raised foot onto the plate, the longest uninterrupted period of single-legged stance was extracted. Subsequently, the Centre-of-Pressure (CoP) position was determined from the averaged pressure distribution during each data frame for each participant’s foot. All data processing was conducted in MATLAB 2022b (Mathworks, Natwick, USA). CoP position time series were low-pass filtered at 10 Hz, and the displacement in each direction was used to calculate a frame-by-frame position change vector in the horizontal plane to yield a direction-unspecific rate of change measure of body sway (dCoP). Within-trial body sway variability was defined as the standard deviation of the rate of change measure (SD dCoP). Absolute and percentage changes in body sway variability resulting from the availability of interpersonal contact were calculated. Figure S1 provides illustrative data traces of a pair of individuals standing with eyes closed with and without IPT.

### Statistical analysis

For the analysis of the influencing factors on the relative sway change during interpersonal touch, we first excluded individuals as outliers with respect to age, anthropometric parameters (height, weight, BMI), an individual’s balancing skill in single-legged stance and relative interindividual differences in balancing skills, as well as the paired partner of the outlier individual. We first computed a 2×2 repeated measures ANOVA to observe the commonly observed effects of vision and IPT on body sway. To investigate if the variability/stability of an individual’s balancing skill shows an inverse or quadratic relationship with age (4, 5) and a quadratic relationship with BMI, we computed a curved fitting analysis (Fig. S5). Curve fitting was repeated based on age and BMI also for relative benefit of IPT (relative change in stability of balancing skills due to IPT) (Fig. S2).

To find the number of personalised performance clusters present within the dataset, based on individual responses to IPT (relative sway change), personal characteristics, as well as the partner’s characteristics, we performed a hierarchical cluster analysis (SPSS 28.0.1.0, IBM SPSS Statistics, Ward’s method). The number of performance clusters and participants’ assignments were then used as a moderator in the next step of backward bootstrapped regression analysis to predict the benefit of IPT (relative sway change) with an individual’s balancing skill in single-legged stance (SD dCoP, no IPT), interindividual difference in balancing skill, sex, motor developmental potential (age mean-centred inverse; assumption of benefit of IPT approaching an asymptote in children), extreme BMI (BMI mean-centred squared; assumption of benefit of IPT increasing in extreme BMI). Further, relative differences between interaction partners in sex, age and BMI were included (Fig. 1). As we expected the effect of an individual’s balancing skill and interindividual differences in balancing skills on the benefit of IPT to be moderated by personalised performance cluster assignment, age, sex and BMI, for which we included the interaction parameters. For this, we multiplied an individual’s balancing skill (SD dCoP, no IPT) and relative interindividual differences in balancing skills with the following moderators: motor developmental potential, and BMI mean-centred squared, as well as dummy coded cluster assignment and sex). To further validate the expected serially mediated relationships of age and anthropometric factors with individual balancing skills, as well as between differences in age and anthropometric factors on the interindividual differences in balancing skills, in the next step, we computed a bootstrapped serial mediation analysis (N=1000, 95%CI, seed 2021) (PROCESS v4.0; 60). The results of the relationship of age and anthropometric factors with an individual’s balancing skills and the benefit of IPT and detailed results of the mediation analysis can be found in the supporting materials. Bootstrapping was applied to augment data, as each condition consisted only of a single trial for every participant, as well as to counteract a potential non-normal distribution of the data. All statistical analyses were performed (SPSS 28.0.1.0, IBM SPSS Statistics). The significance level was set to 0.05, and the statistical tendency level to 0.10. Cohen’s d and partial η2 are reported as effect sizes with low, moderate and strong effect defined as d=0.2, d=0.5, d=0.8 and η2=0.01, η2=0.06 and η2=0.14, respectively.

## Results

One hundred and forty-four individuals in the age range from 4 to 63 years (70 f, 74 m) were included in the analysis. Individuals were generally more stable with eyes open and when IPT was available (Fig. 2, Table S.1). Further, the benefit of IPT was greater in the Eyes closed (EC) condition (Mdiff=75.73 (36%), p<0.001, 95%CI [57.17 94.30]) than in the Eyes open (EO) condition (Mdiff=13.58 (23%), p<0.001, 95%CI [9.00 18.16]; F(1,143)=44.72, p<0.001, η2=0.24). The bootstrapped descriptive statistics of the whole group are further shown in Tables S1 and S2.

**Figure 2.**
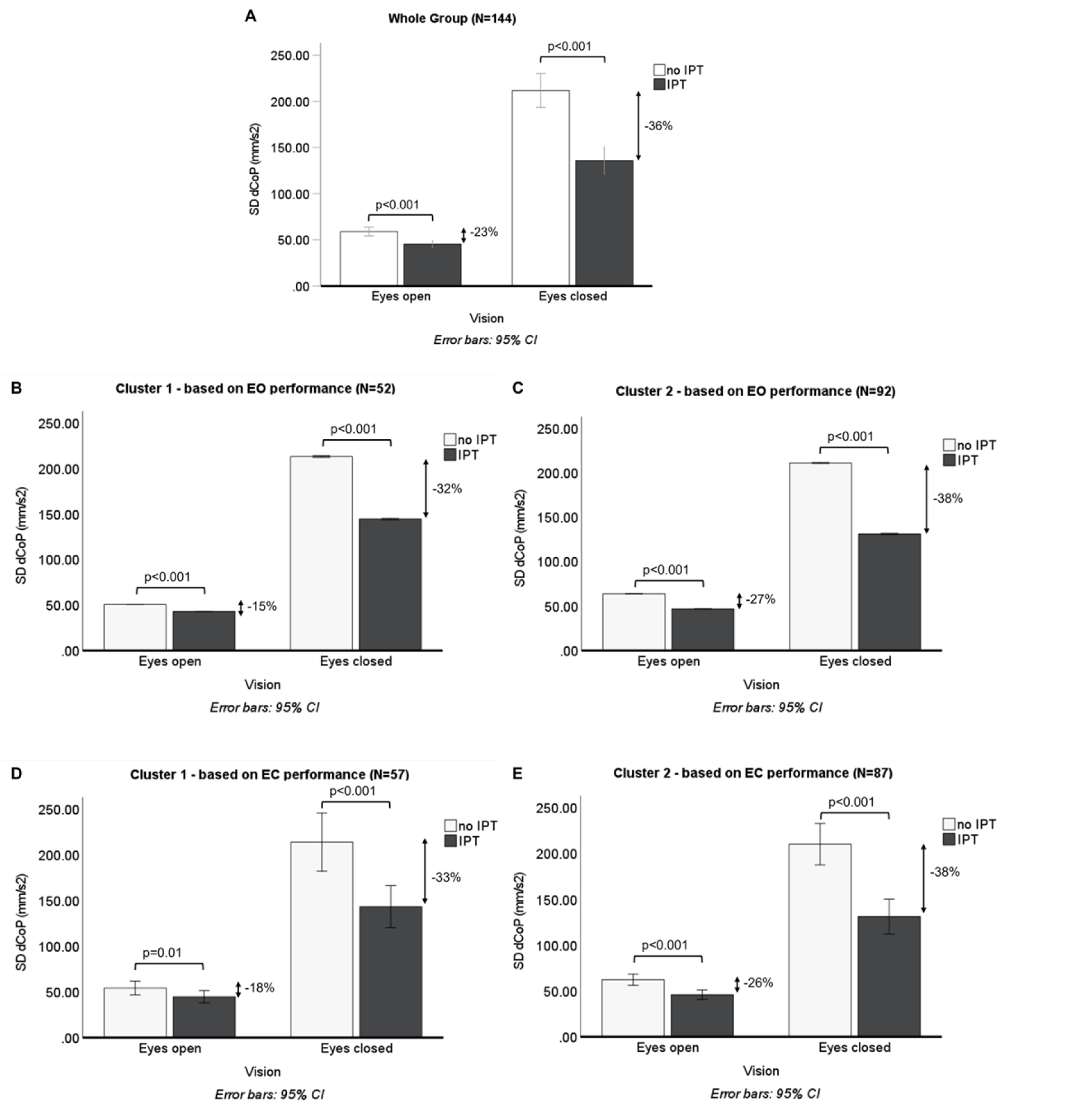
Bar graphs of the effect of vision and interpersonal tactile interaction on variability of body sway velocity (marginal means) for the entire participants sample (A) and by performance cluster assignments based on behaviour during Eyes open condition (B, C) and Eyes closed condition (D, E). In general, a greater benefit of IPT is observable on the Eyes closed condition and for the participants assigned to Cluster 2 (‘vulnerable’ participants). Cluster 1: more skilled, older, taller and heavier participants; Cluster 2: more vulnerable participants. EO: Eyes open, EC: Eyes closed; IPT: interpersonal touch.

A hierarchical cluster analysis, considering individuals’ sway change due to the presence of IPT relative to standing without IPT, personal characteristics and relative characteristics compared to the interaction partner, revealed two main clusters of participants (EO condition: Cluster 1: N=52, Cluster 2: N=92; EC condition: Cluster 1: N=57, Cluster 2: N=87). Participants’ demographics and individual characteristics for the two clusters in the EO and EC conditions without IPT are shown in Tables A3 and A4, respectively. Comparing the two clusters, it becomes apparent that the first cluster included older, taller and heavier individuals, while the second cluster consisted of younger, smaller and lighter individuals, and consequently with a lower BMI.

Furthermore, in the EO condition, individuals in the second cluster demonstrated reduced single-legged balancing skill (in terms of greater variability of CoP velocity), relatively lower balancing skill within a pair, and a greater benefit of IPT (greater reduction of variability of CoP velocity due to IPT) compared to individuals in the first cluster. These differences between the clusters, however, were only observed in the EO, but not in the EC condition, which indicated that lack of vision also challenged the individuals in the first cluster considerably.

Figure 3 depicts the relationship between interindividual differences in age-related motor experience and the benefit of IPT for each performance cluster separately. This figure further distinguishes between individuals whose partner was assigned to the same or to a different cluster. For the EO condition (Fig. 3. A, B), differences in the benefit of IPT between the same and different partner cluster assignments were not observed, neither for the first nor for the second cluster (bootstrapped t-tests: Cluster One: same cluster partner (N=20): Mean=-8.58, BCa95% [−19.42 3.63], SD=27.38, BCa95%CI [10.30 37.97]; different cluster partner (N=32): Mean=-6.97, BCa95%CI [−14.18 1.35], SD=24.58, BCa95%CI [11.74 32.88]; Mdiff=1.62, p=0.847, BCa95%CI [−13.24 15.47], Cohen’s d=0.06 95CI [−13.24 15.47]; Cluster Two: same cluster partner (N=60): Mean=-14.29, BCa95%CI [−22.03 −6.85], SD=30.30, BCa95%CI [22.77 37.05]; different cluster partner (N=32): Mean=-21.99, BCa95%CI [−30.85 - 13.64], SD=25.00, BCa95%CI [17.55 30.56], Mdiff=-7.70, p=0.193, BCa95%CI [−18.73 2.82], Cohen’s d=-0.27 95CI [−0.70 0.16]).

**Figure 3.**
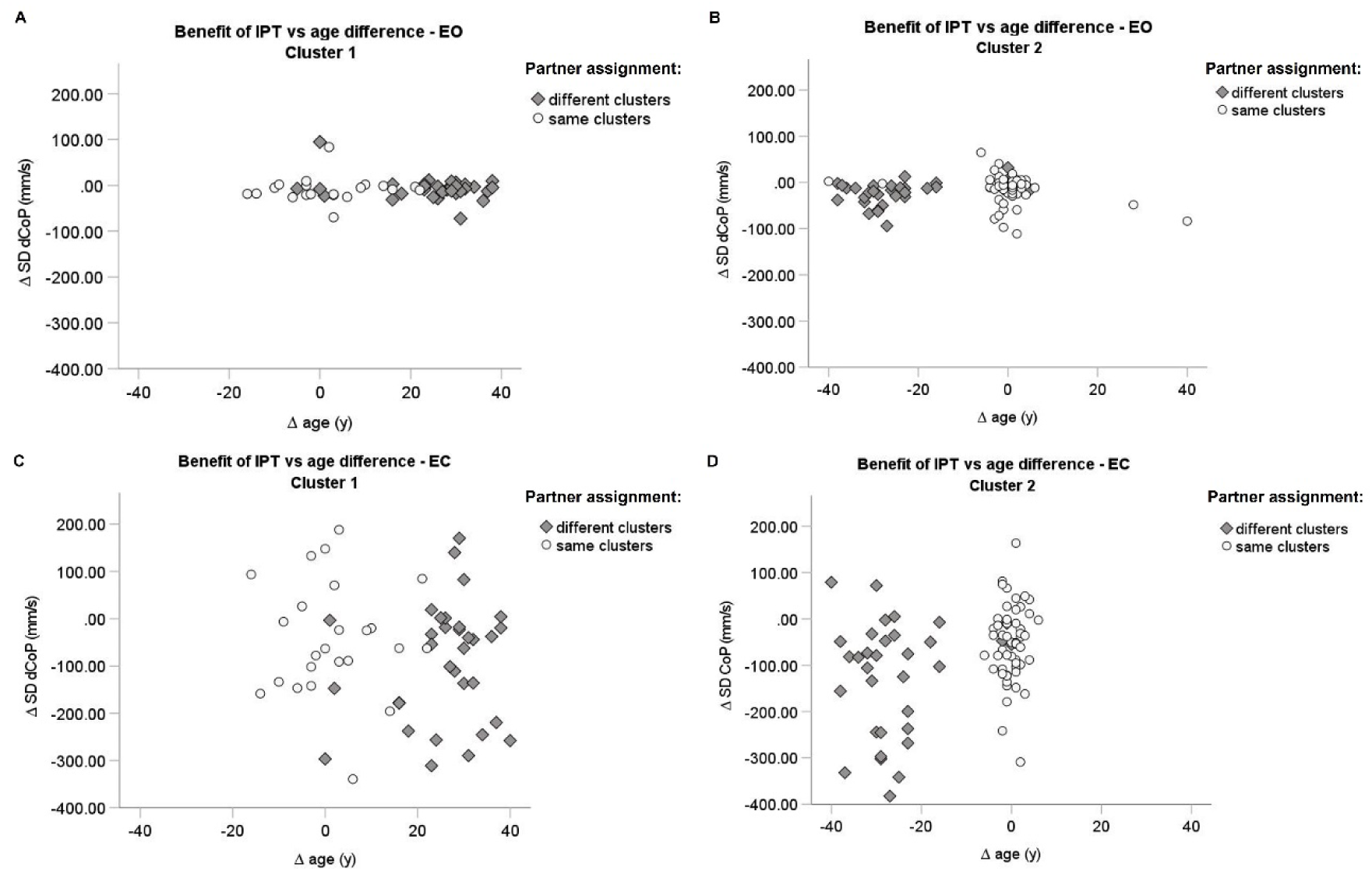
Scatter plot of the benefit of IPT in the Eyes open condition (A, B) and Eyes closed condition (Fig. 3. C, D) by cluster assignment of an individual and cluster assignment of their interaction partner. Cluster 1: more skilled, older, taller and heavier participants; Cluster 2: more vulnerable participants. EO: Eyes open, EC: Eyes closed; IPT: interpersonal touch.

In contrast, in the EC condition, for the individuals in the second cluster a significant difference between individuals whose partner was assigned to the same vs. different cluster was observed. An individual with a partner in the other (first) cluster showed a greater sway reduction (N=33: Mean=-122.60, BCa95%CI [−166.18 −76.14], SD=122.72, BCa95% [98.49 139.67]) compared to individuals with a partner assigned to the same (second) cluster (N=54: Mean=-52.54, BCa95%CI [−75.77 −30.80], SD=81.84, BCa95%CI [64.72 96.03]; Mdiff=-70.05, p=0.007, Bca95%CI [−118.53 −23.45], Cohen’s d=-0.71 95CI [−1.15 −0.26]). However, comparing individuals in both clusters directly against each other as a function of the partner assignments showed no differences between clusters, neither for partner assignment to the same nor to different clusters (Mdiff=11.26, p=0.670, BCa95%CI [−43.18 64.08], Cohen’s d=0,12 95CI [−0.36 0.60]; Mdiff=30.72, p=0.335, BCa95%CI [−28.68 91.87], Cohen’s d=0.25 95CI [−0.24 0.73]), respectively.

Bootstrapped regression analysis indicated that an individual’s amount of relative sway change could be explained to 46% (p<0.001) in the EO condition and to 52% (p<0.001) in the EC condition (Fig. 4 A and B, respectively). Both conditions had in common, that individuals in the second performance cluster benefitted more from an older, thus more experienced, interaction partner. (EO: B=0.36, Bias=-0.03, SE=0.11, p=0.002, BCa95% [0.15 0.57], β =0.17; EC: B=1.44, Bias=0.00, SE=0.57, p=0.013, Bca95% [0.25 2.54], β =1.53).

**Figure 4.**
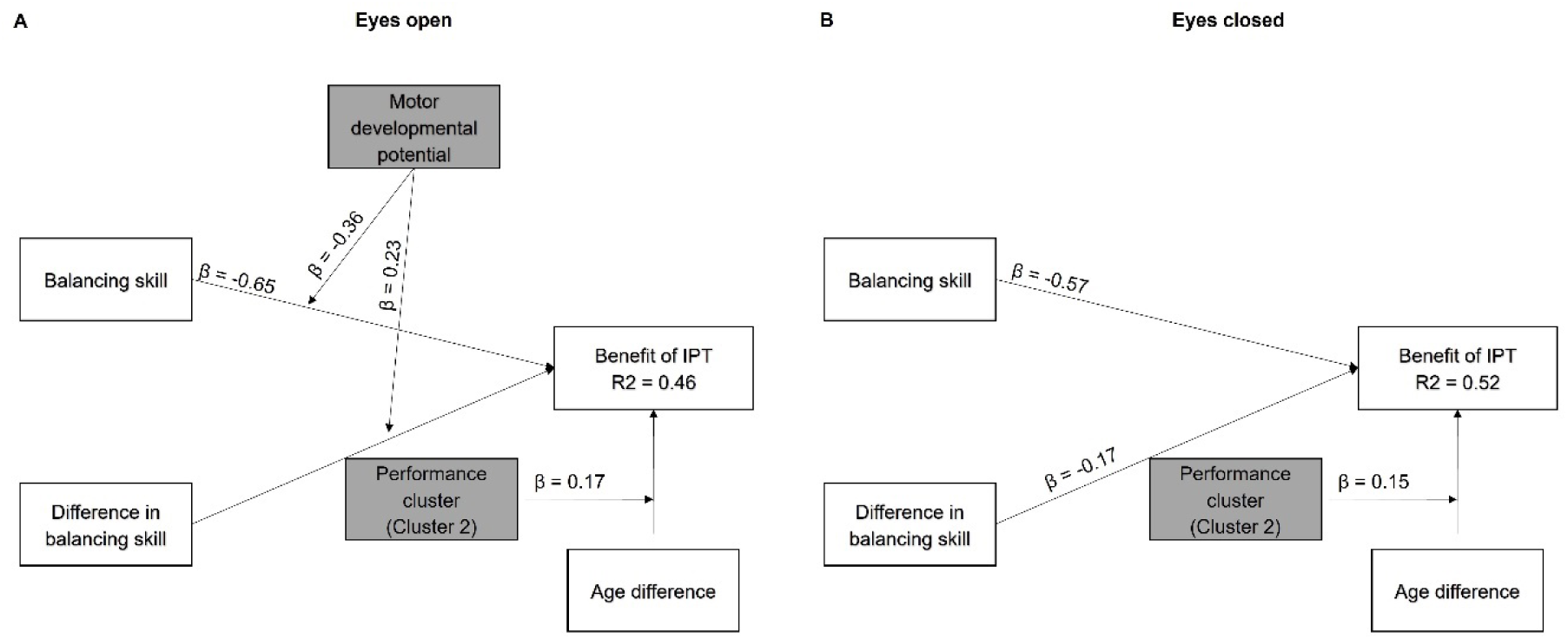
Statistical model of significant influential factors on the benefit of IPT. (A) In the Eyes open condition, the benefit of IPT depends on 1) an individual’s balancing skill, 2) an interaction between the balancing skill by motor developmental potential, 3) an interaction between the interindividual differences in balancing skills and motor developmental potential, and 4) the age differences for individuals in Cluster 2. (B) In the Eyes closed condition, the benefit of IPT depends on 1) an individual’s balancing skill and the 2) interindividual difference in balancing skills and 3) the age differences for individuals in Cluster 2. IPT: interpersonal touch, β: standardized coefficient.

Nevertheless, differences between regression models for the EO and EC conditions were also found. Although in both visual conditions, individuals with less balancing skills (greater sway variability) showed an increased benefit of IPT (greater relative reduction in sway variability) (EO: B=-0.63, Bias=0.06, SE=0.09, p<0.001, BCa95% [−0.80 −0.42], β=-0.65; EC: B=-0.57, Bias=0.03, SE=0.09, p<0.001, BCa95% [−0.74 −0.42], β=-0.57), in the EO condition the influence of balancing skills was greater in individuals with a greater developmental potential (younger age with greater motor developmental potential (B=-0.64, Bias=0.04, SE=0.32, p=0.025, Bca95% [−1.09 0.03], β=-0.36).

Moreover, during EO, differences in balancing skills influenced more the benefit of IPT in individuals with a greater motor developmental potential, and thus, younger individuals (B=43.65, Bias=-10.13, SE=26.74, p=0.082, BCa95% [1.49 67.40], β=0.23). It means that the influence of interindividual differences in balancing skill as well as the influence of an individual’s balancing skill on the benefit of IPT was enhanced with younger age (Fig. S4). In contrast, in the EC condition the influence of an individual’s balancing skill and of interindividual differences in balancing skills was not moderated by an individual’s motor developmental potential but revealed, in general, a greater benefit of IPT for less skilled individuals as reported above, and for individuals paired with a more skilled interaction partner (B=-21.77, Bias=0.51, SE=8.70, p=0.016, Bca95% [−38.03 −1.66], β =-0.17).

An additional serial regression analysis was performed which showed that the variance of an individual’s balancing skill could be explained by anthropometry and age to 14% and 18% only in both visual feedback conditions (Fig. S6).

## Discussion

Our study assessed the extent to which an individual’s characteristics such as current age and potential in motor development (age inverse), height and weight of the body (via BMI), and an individual’s balancing skill (in terms of body sway during single-legged stance without support) in combination with contextual factors such as the relative differences in the personal characteristics between interaction partners determine the stabilising benefit of the interpersonal touch (IPT).

We confirmed that IPT improves stability of balance in single-legged stance (reduced variability in balancing performance) and that stance stabilisation is greater with IPT when visual feedback is not available. IPT resulted in sway reductions of 15-23% with and 32-38% without vision. This is a greater effect compared to previously observed reductions of 9-18% due to IPT in normal bipedal and Tandem-Romberg stances (greater reduction in Tandem-Romberg stance; 13, 61). The augmented reductions that we observed may be a consequence of the more unstable nature of a single-legged stance compared to previous studies with more stable stance postures. The amount of reduction in variability in balancing performance, especially without vision, is comparable to the effect of touching a static surface (20-31%) (62, 63) when grasping another individual’s shoulder (37%; 20), or when touching an artificial stabilising interaction partner (34-36%; 19).

Our findings indicate that the benefit of IPT is not driven by an individual’s anthropometry or any interindividual differences in anthropometry. In our present study, we found that an individual’s stabilisation benefit from IPT was affected by an individual’s balancing skill and by the relative differences between interaction partners’ individual balancing skills as well as age differences instead. The amount of explained variance of balance performance by anthropometry that we observed was comparable to the explained variance reported in previous studies (3, 4, 7, 64). Thus, the influence of anthropometry on an individual’s balancing skill and, thus, on the benefit of IPT are rather minor. This observation confirms a meta-analysis by Schmuckler (28), who found an anthropometric factor such as body height to influence balance stability in children only slightly. In the age range from 3 to 7 years body balance improved with increasing body height, possibly reflecting maturing sensorimotor control of balance, while at greater age body height imposed biomechanical constraints (2, 65).

Irrespective of the visual feedback condition, IPT benefit was greater for more vulnerable, generally younger, smaller, and lighter, individuals when interacting with a relatively older interaction partner compared to themselves. This was observed although these participants were no longer in infancy or early childhood and therefore not reliant on any external, interpersonal locomotor support any longer. At an average age of around 11 years the more vulnerable participants of the second cluster supposedly have mastered their fundamental motor skills. Furthermore, as expected, a person’s lower level of balancing skill was associated with greater balance improvements when IPT was available, which was observed independent of motor developmental potential when balance stability was challenged by the lack of visual feedback. On the other hand, when visual feedback was available. the individual balancing skill had a greater influence on the benefit of IPT for individuals with a greater developmental potential, and thus, younger individuals,

Remarkably, we also observed that individuals, who were relatively more unstable compared to their interaction partner during measurements without IPT, showed a greater benefit from IPT without differences in their actual standing postures, which emphasises the importance of their balancing skills relative to their partners balancing skills. This observation parallels previous reports where more unstable individuals standing in a Tandem-Romberg stance benefit more from IPT than their interaction partners standing in a more stable normal bipedal stance (61). While this was observable independent of motor developmental potential when visual feedback was not available, in the condition with visual feedback individuals a relatively more stable interaction partner was more relevant for individuals with a greater motor developmental potential, and thus younger individuals.

The moderation of the effect of both balancing skills and differences in balancing skills on the benefit of IPT by an individual’s motor developmental potential implies that the motor developmental potential of a participant (meaning balance control being refined less by experience) impacts on the processing of visual information for balance control. In this situation, the more vulnerable individuals demonstrated comparatively high intra- (and inter-) individual variability in their level of balancing skill and the relative difference to their interaction partner (Fig. 2, Supporting Tables S3 and S4, Figs. S2 - S4). In his meta-analysis, Schmuckler (28) rested the conclusion that the influence of proprioceptive information dominates balance control in children on the observed variable effects of stance width. This conclusion needs to be qualified by the fact that different stance postures not only entail an altered configuration of proprioceptive input but also confound differences in the intrinsic stability of a given stance posture (a wider stance is more stable in the mediolateral direction at least). Nevertheless, a single-legged stance as chosen in our present study generates quite salient muscle activations and proprioceptive feedback so that we do not see a contradiction with Schmuckler’s conclusions (28).

The efficacy of balance control during a challenging single-legged stance may be limited by immature, less efficient multisensory integration, and less refined internal body representations in children and adolescents compared to adults (38). Thus, a stronger influence of interpersonal differences in balancing skill during IPT may be associated with a greater susceptibility of vulnerable individuals to the haptic feedback received in terms of the interaction force (25). The processing of proprioception for balance control remains underdeveloped until around 9 years of age (26). Multisensory reweighting and integration continue to develop even in late childhood and adolescence (2, 66, 67). Moreover, children have been shown to rely more on sensory feedback compared to adults (36, 68) and they reweight and integrate inter-modal sensory information less adequately. For example, children are less capable to uncouple from tactile feedback at destabilising frequencies (25, 69) and are more responsive to a wider range of tactile stimulus frequencies. Similarly, as in older age (42), noisier sensorimotor processing (measurement noise, estimator/computational noise, process/command noise), and a less accurate and precise, and thus more uncertain internal representations may require recalibration and refinement based on multisensory feedback during development. Thus, it is not surprising that these more vulnerable individuals showed greater intra- and interpersonal variability (Figs. S2 and S4) and a greater reliance on an older and more stable interaction partner than less vulnerable individuals and individuals with less motor developmental potential (Fig. 3).

It is remarkable, however, that a similar moderating effect of motor developmental potential was not observed without visual feedback. In the condition without visual feedback, an individual’s balancing skill and any interpersonal differences in balancing skills exerted a comparable influence on stability across the entire age range of the study. As standing on one leg without visual feedback is also challenging for adults, postural responses to IPT may also become more variable in adults (Fig. 2, Supporting Table S4, Figs. S2 and S4). Nevertheless, the vulnerable individuals with a partner from the other cluster showed greater sway reduction with IPT with eyes closed than those vulnerable individuals whose interaction partner was equally vulnerable (Fig. 4). A similar difference in IPT sway reduction between non-vulnerable individuals with partners from the same cluster or partners from the other cluster was not observed.

The apparently greater dependency on an interaction partner with more dissimilar personal characteristics may indicate that more vulnerable participants, such as children and adolescents, possess less precise representations of their own body and its movement dynamics. Such limitations may become especially relevant when the visual channel does not contribute to feedback control of body balance. Uncertainty about the sensory consequences of own balance adjustments may make it harder to distinguish between self-induced sensory consequences and consequences of an interaction partner’s balance adjustments. Therefore, when the relative differences to the characteristics of the interaction partner are more pronounced, a distinction between self- and other-induced sensory feedback may be facilitated, especially when the partner is more stable. Better distinction between self-evoked sway dynamics and dynamics evoked by the interaction partner might lead to more precise balance adjustments.

Moreover, when the representation of oneself is less precise and one’s stance is unstable, individuals may be more likely to confuse their movement consequences with the other’s. This confusion may lead to the commonly known interpersonal entrainment and resonance effects (e.g. synchronisation of each other’s movements in the same direction). Lower intraindividual variability of a more mature interaction partner may increase the predictability of the partner (70) and therefore raise the likelihood of distinguishing the sources of sensory feedback for the more vulnerable partner during IPT. In contrast, if more mature individuals are affected less by sensory and control system noise and possess more efficient multisensory integration processes as well as more accurate and precise internal representations, they may be able to rely more on their own proprioceptive feedback and be less responsive to the haptic feedback granted by IPT. Their internal representations may allow a greater degree of predictive control when interacting with a more unstable or unreliable interaction partner, so any dependency on any partner’s characteristics is diminished, which nevertheless does not preclude a beneficial utilisation of IPT.

Future studies are required to gain more insights into the role of own and relative partner characteristics on the benefit of IPT in the old (>80y) or oldest (>90y) population. As older aged individuals are known to rely more on concurrent delayed and more noisy multisensory feedback for continuously updating internal models, we hypothesise to find similar effects in older age; however, in the opposite direction, meaning that older individuals are expected to benefit more from a relatively younger interaction partner.

## Conclusions

This study showed that anthropometrical parameters, such as body hight and weight, only play a minor role for the benefit of interpersonal touch, and that the developmental motor experience as well the single-legged balancing skill play major roles. Besides an individual’s characteristics, such as balancing skill and age-related motor experience, also the relative differences in characteristics to an interaction partner contributed to the benefit of interpersonal touch on balance stability. We found that especially children and individuals with reduced balancing skills benefited more from interpersonal touch and that these individuals also benefited more from interacting with a relatively older partner. IPT is a balance support strategy, that seems to benefit stability in a broad range of individuals, especially vulnerable individuals interacting with more stable partners, and in challenging stance contexts, where mutual destabilization is likely. Thus, IPT is a promising approach in clinical contexts, such as routine activities and training sessions.

## Author contributions

L.J. has designed and planned the primary study. L.J. has collected the data, K.S., L.J. and D.L. have designed the secondary analysis, L.J. and K.S. have processed the data, K.S., has analysed the data. K.S., L.J., have prepared the manuscript, L.J. and D.L. have supervised.

## Data availability statement

All extracted data files are available from the figshare database (10.6084/m9.figshare.27827127).

## Supporting information and materials

**Table S1.** Descriptive statistics of the entire participant sample: individual characteristics. EO: Eyes open, EC: Eyes closed; IPT: interpersonal touch.

| Parameters |  |
| --- | --- |
| Sex: f/m | N=70 (48.6%) / N=74 (51.4%) |
| Age-related motor experience (y) | M=20.65, BCa95% [18.32 22.97]<br>SD=15.25, BCa95% [13.89 16.43]<br>Min=4, Max=63 |
| Height (m) | M=1.54, BCa95% [1.51 1.57]<br>SD=0.20, BCa95% [0.18 0.21]<br>Min=1.12, Max=1.94 |
| Weight (kg) | M=49.87, BCa95% [46.10 53.55]<br>SD=24.61, BCa95% [22.31 26.91]<br>Min=12, Max=123 |
| BMI (kg/m<sup>2</sup>) | M=19.41, BCa95% [18.56 20.31]<br>SD=5.49, BCa95% [4.97 5.99]<br>Min=9.3 , Max=35.5 |
| Balancing skill, EO - no IPT (SD dCoP; mm/s<sup>2</sup>) | M=59.07, BCa95% [54.76 63.98]<br>SD=28.62, BCa95% [23.55 33.22]<br>Min=19.60, Max=167.16 |
| Balancing skill, EC - no IPT (SD dCoP; mm/s<sup>2</sup>) | M=211.74, BCa95% [192.25 230.48]<br>SD=111.49, BCa95% [99.85 122.01]<br>Min=51.47, Max=513.642 |
| Pairing: Same sex / different sex | N=76 (52.8%), N=68 (47.2%) |

**Table S2.** Descriptive statistics of the entire participant sample: relative interindividual differences in characteristics between interaction partners. EO: Eyes open, EC: Eyes closed; IPT: interpersonal touch.

|  |  |
| --- | --- |
| Difference in age-related motor experience (y) | M=0.30, BCa95% [-2.84 3.48]<br>SD=19.30, BCa95% [17.57 20.87]<br>Min=-40, Max=40 |
| Height difference (m) | M=0.00, BCa95% [-0.04 0.04]<br>SD=0.23, BCa95% [0.20 0.25]<br>Min=-67, Max=67 |
| Weight difference (kg) | M=0.00, BCa95% [-4.73 5.16]<br>SD=31.07, BCa95% [27.58 34.35]<br>Min=-78, Max=78 |
| BMI difference (kg/m<sup>2</sup>) | M=0.00, BCa95% [-1.16 1.21]<br>SD=7.32, BCa95% [6.59 7.99]<br>Min=-16.20, Max=16.20 |
| Difference in balancing skills, EO (mm/s<sup>2</sup>) | M=0.16, BCa95% [0.04 0.29]<br>SD=0.71, BCa95% [0.51 0.91]<br>Min=-0.81, Max=4.33 |
| Relative benefit of IPT, EO (relative sway change due to IPT EO;mm/s<sup>2</sup>) | M=-13.58, BCa95% [-18.41 -9.00]<br>SD=27.83, BCa95% [22.63 32.77]<br>Min=-111.18, Max=94.87 |
| Percentage benefit of IPTm EO (percentage sway change due to IPT, EO (%)) | M=-17.01, BCa95% [-22.32 -10.93]<br>SD=34.14, BCa95% [22.61 41.79]<br>Min=-79.89, Max=153.31 |
| Difference in balancing skills, EC (mm/s<sup>2</sup>) | M=0.26, BCa95% [0.12 0.42]<br>SD=0.89, BCa95% [0.50 0.91]<br>Min=-0.81, Max=4.33 |
| Relative benefit of IPT, EC (relative sway change due to IPT, EC (mm/s<sup>2</sup>)) | M=-75.74, BCa95% [-94.93 -56.53]<br>SD=112.71, BCa95% [99.96 124.24]<br>Min=-382.65, Max=188.40 |
| Percentage benefit of IPT, EC (sway change due to IPT, EC (%)) | M=-24.25, BCa95% [-32.17 -15.53]<br>SD=52.90, BCa95% [42.47 62.39]<br>Min=-85.27, Max=216.08 |

**Table S3.**
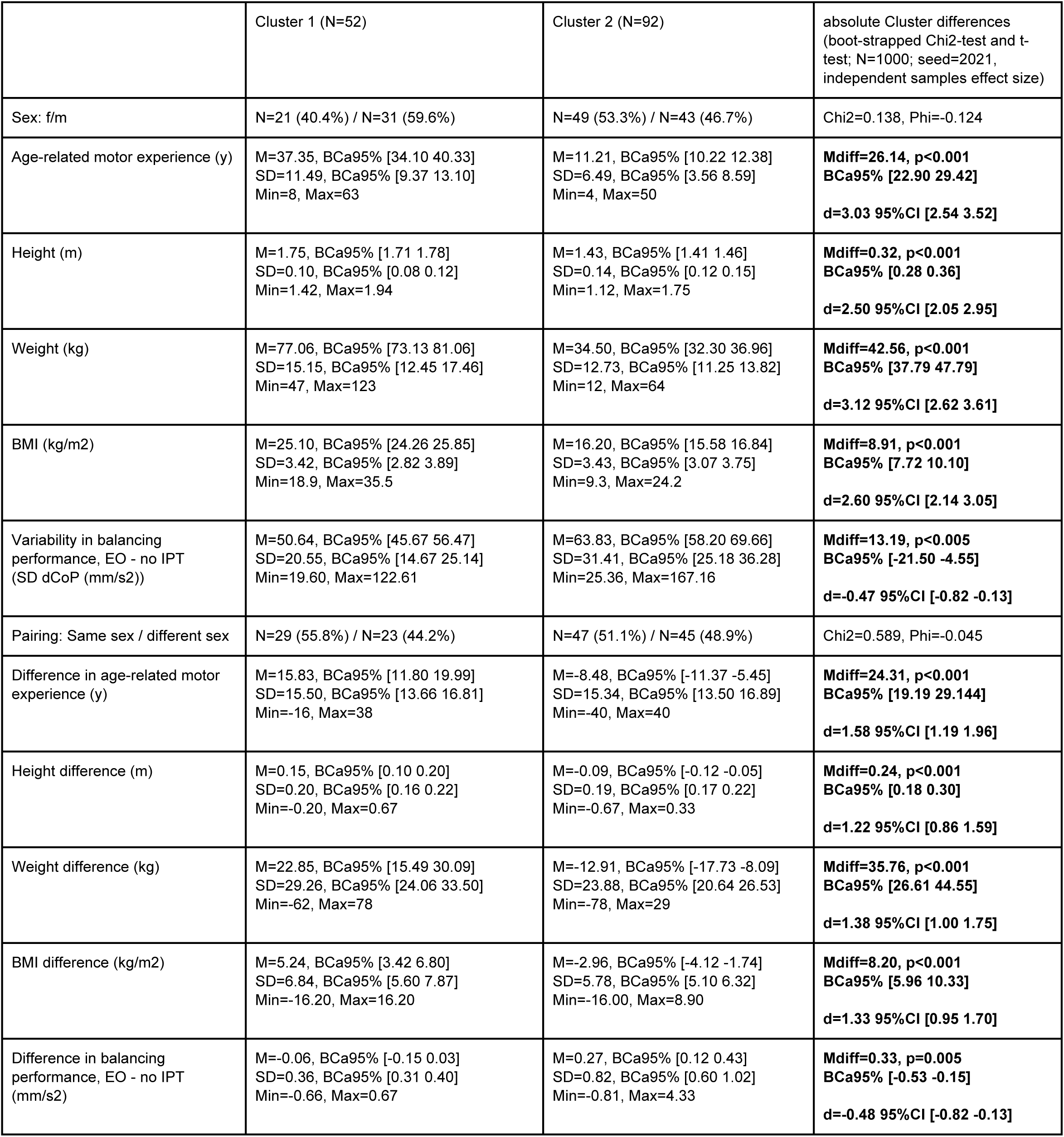
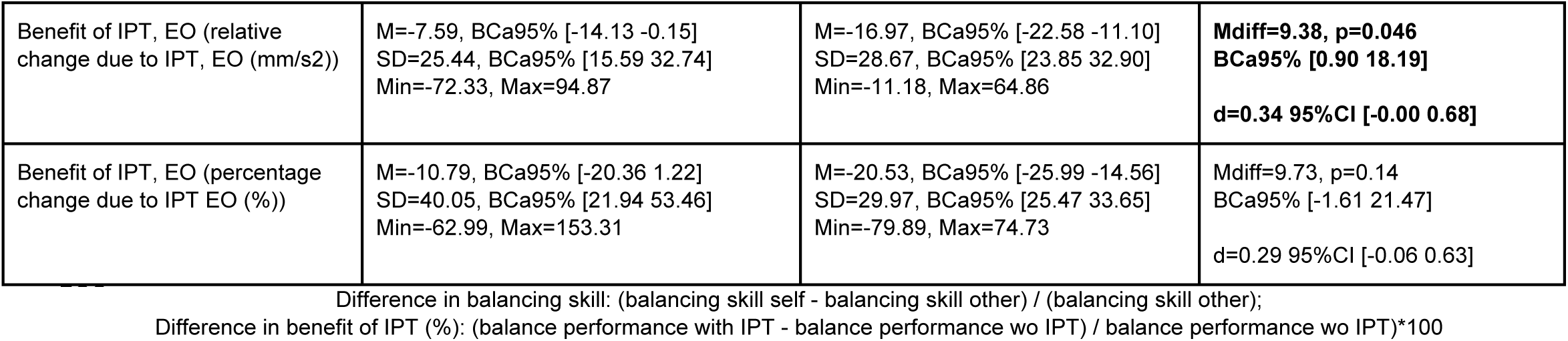
Participant characteristics for the two performance clusters based on the Eyes open condition. Significant cluster differences are indicated by a star and statistics are shown in the last column (equal variances not assumed). EO: Eyes open, EC: Eyes closed; IPT: interpersonal touch.

|  | Cluster 1 (N=52) | Cluster 2 (N=92) | absolute Cluster differences<br>(boot-strapped Chi2-test and t-<br>test; N=1000; seed=2021,<br>independent samples effect size) |
| --- | --- | --- | --- |
| Sex: f/m | N=21 (40.4%) / N=31 (59.6%) | N=49 (53.3%) / N=43 (46.7%) | Chi2=0.138, Phi=-0.124 |
| Age-related motor experience (y) | M=37.35, BCa95% [34.10 40.33]<br>SD=11.49, BCa95% [9.37 13.10]<br>Min=8, Max=63 | M=11.21, BCa95% [10.22 12.38]<br>SD=6.49, BCa95% [3.56 8.59]<br>Min=4, Max=50 | **Mdiff=26.14, p<0.001**<br>**BCa95% [22.90 29.42]**<br>**d=3.03 95%CI [2.54 3.52]** |
| Height (m) | M=1.75, BCa95% [1.71 1.78]<br>SD=0.10, BCa95% [0.08 0.12]<br>Min=1.42, Max=1.94 | M=1.43, BCa95% [1.41 1.46]<br>SD=0.14, BCa95% [0.12 0.15]<br>Min=1.12, Max=1.75 | **Mdiff=0.32, p<0.001**<br>**BCa95% [0.28 0.36]**<br>**d=2.50 95%CI [2.05 2.95]** |
| Weight (kg) | M=77.06, BCa95% [73.13 81.06]<br>SD=15.15, BCa95% [12.45 17.46]<br>Min=47, Max=123 | M=34.50, BCa95% [32.30 36.96]<br>SD=12.73, BCa95% [11.25 13.82]<br>Min=12, Max=64 | **Mdiff=42.56, p<0.001**<br>**BCa95% [37.79 47.79]**<br>**d=3.12 95%CI [2.62 3.61]** |
| BMI (kg/m2) | M=25.10, BCa95% [24.26 25.85]<br>SD=3.42, BCa95% [2.82 3.89]<br>Min=18.9, Max=35.5 | M=16.20, BCa95% [15.58 16.84]<br>SD=3.43, BCa95% [3.07 3.75]<br>Min=9.3, Max=24.2 | **Mdiff=8.91, p<0.001**<br>**BCa95% [7.72 10.10]**<br>**d=2.60 95%CI [2.14 3.05]** |
| Variability in balancing<br>performance, EO - no IPT<br>(SD dCoP (mm/s2)) | M=50.64, BCa95% [45.67 56.47]<br>SD=20.55, BCa95% [14.67 25.14]<br>Min=19.60, Max=122.61 | M=63.83, BCa95% [58.20 69.66]<br>SD=31.41, BCa95% [25.18 36.28]<br>Min=25.36, Max=167.16 | **Mdiff=13.19, p<0.005**<br>**BCa95% [-21.50 -4.55]**<br>**d=-0.47 95%CI [-0.82 -0.13]** |
| Pairing: Same sex / different sex | N=29 (55.8%) / N=23 (44.2%) | N=47 (51.1%) / N=45 (48.9%) | Chi2=0.589, Phi=-0.045 |
| Difference in age-related motor<br>experience (y) | M=15.83, BCa95% [11.80 19.99]<br>SD=15.50, BCa95% [13.66 16.81]<br>Min=-16, Max=38 | M=-8.48, BCa95% [-11.37 -5.45]<br>SD=15.34, BCa95% [13.50 16.89]<br>Min=-40, Max=40 | **Mdiff=24.31, p<0.001**<br>**BCa95% [19.19 29.144]**<br>**d=1.58 95%CI [1.19 1.96]** |
| Height difference (m) | M=0.15, BCa95% [0.10 0.20]<br>SD=0.20, BCa95% [0.16 0.22]<br>Min=-0.20, Max=0.67 | M=-0.09, BCa95% [-0.12 -0.05]<br>SD=0.19, BCa95% [0.17 0.22]<br>Min=-0.67, Max=0.33 | **Mdiff=0.24, p<0.001**<br>**BCa95% [0.18 0.30]**<br>**d=1.22 95%CI [0.86 1.59]** |
| Weight difference (kg) | M=22.85, BCa95% [15.49 30.09]<br>SD=29.26, BCa95% [24.06 33.50]<br>Min=-62, Max=78 | M=-12.91, BCa95% [-17.73 -8.09]<br>SD=23.88, BCa95% [20.64 26.53]<br>Min=-78, Max=29 | **Mdiff=35.76, p<0.001**<br>**BCa95% [26.61 44.55]**<br>**d=1.38 95%CI [1.00 1.75]** |
| BMI difference (kg/m2) | M=5.24, BCa95% [3.42 6.80]<br>SD=6.84, BCa95% [5.60 7.87]<br>Min=-16.20, Max=16.20 | M=-2.96, BCa95% [-4.12 -1.74]<br>SD=5.78, BCa95% [5.10 6.32]<br>Min=-16.00, Max=8.90 | **Mdiff=8.20, p<0.001**<br>**BCa95% [5.96 10.33]**<br>**d=1.33 95%CI [0.95 1.70]** |
| Difference in balancing<br>performance, EO - no IPT<br>(mm/s2) | M=-0.06, BCa95% [-0.15 0.03]<br>SD=0.36, BCa95% [0.31 0.40]<br>Min=-0.66, Max=0.67 | M=0.27, BCa95% [0.12 0.43]<br>SD=0.82, BCa95% [0.60 1.02]<br>Min=-0.81, Max=4.33 | **Mdiff=0.33, p=0.005**<br>**BCa95% [-0.53 -0.15]**<br>**d=-0.48 95%CI [-0.82 -0.13]** |
| Benefit of IPT, EO (relative change due to IPT, EO (mm/s2)) | M=-7.59, BCa95% [-14.13 -0.15]<br>SD=25.44, BCa95% [15.59 32.74]<br>Min=-72.33, Max=94.87 | M=-16.97, BCa95% [-22.58 -11.10]<br>SD=28.67, BCa95% [23.85 32.90]<br>Min=-11.18, Max=64.86 | **Mdiff=9.38, p=0.046**<br>**BCa95% [0.90 18.19]**<br>**d=0.34 95%CI [-0.00 0.68]** |
| Benefit of IPT, EO (percentage change due to IPT EO (%)) | M=-10.79, BCa95% [-20.36 1.22]<br>SD=40.05, BCa95% [21.94 53.46]<br>Min=-62.99, Max=153.31 | M=-20.53, BCa95% [-25.99 -14.56]<br>SD=29.97, BCa95% [25.47 33.65]<br>Min=-79.89, Max=74.73 | Mdiff=9.73, p=0.14<br>BCa95% [-1.61 21.47]<br>d=0.29 95%CI [-0.06 0.63] |

Difference in balancing skill: (balancing skill self - balancing skill other) / (balancing skill other); Difference in benefit of IPT (%): (balance performance with IPT - balance performance wo IPT) / balance performance wo IPT)\*100

**Table S4.**
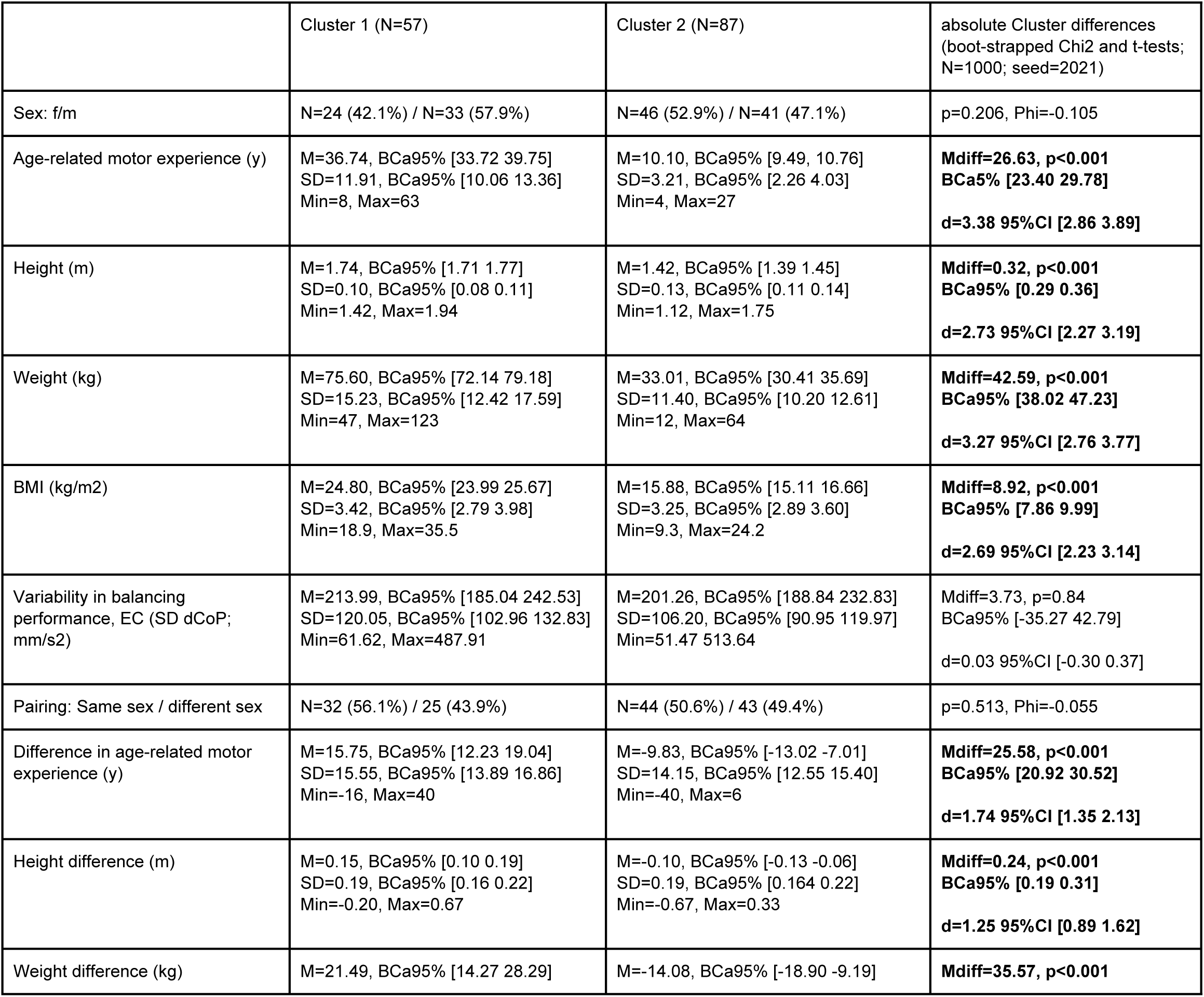
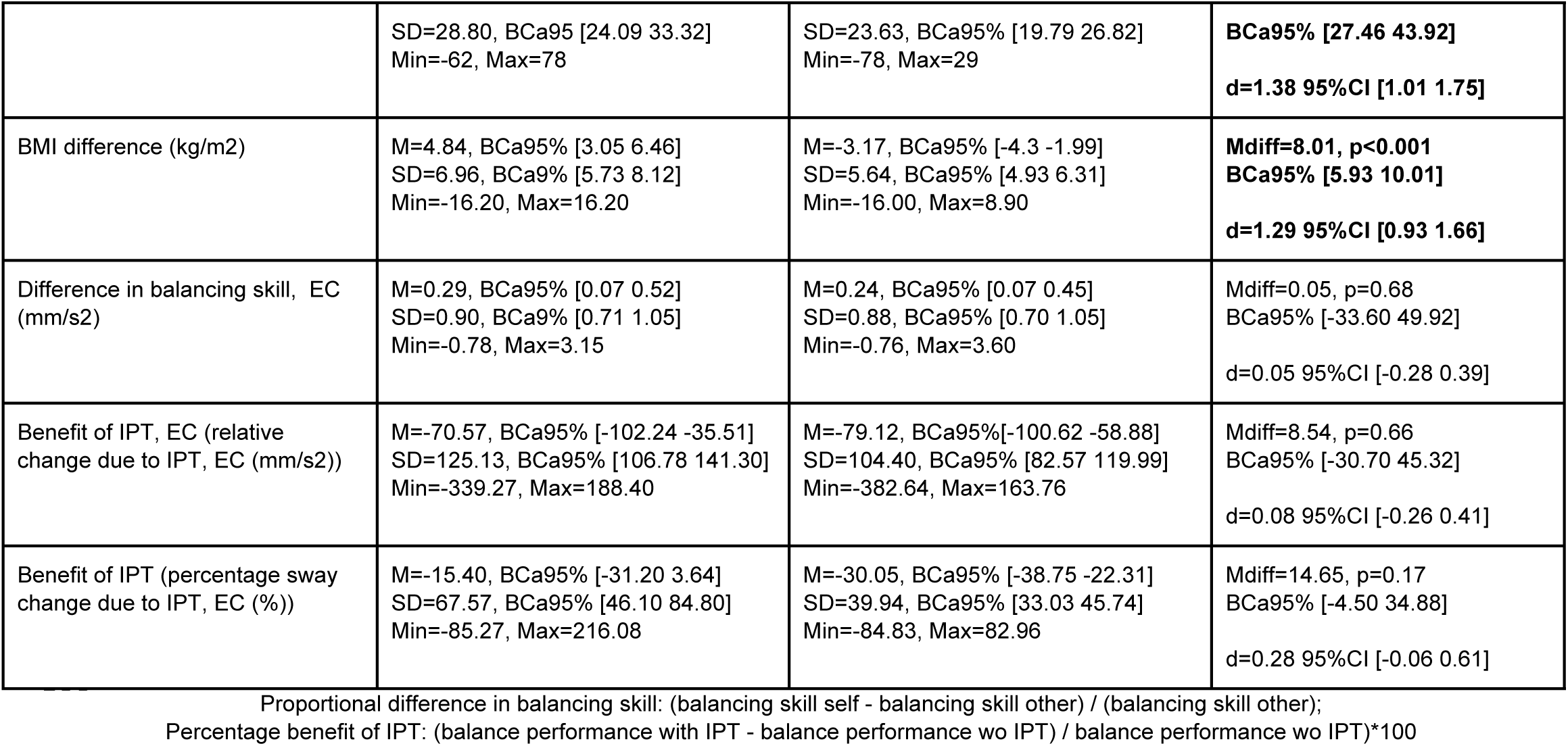
Participant characteristics for the two performance clusters based on Eyes closed condition. Significant cluster differences are indicated by a star and statistics are shown in the last column (equal variances not assumed). EO: Eyes open, EC: Eyes closed; IPT: interpersonal touch.

|  | Cluster 1 (N=57) | Cluster 2 (N=87) | absolute Cluster differences<br>(boot-strapped Chi2 and t-tests;<br>N=1000; seed=2021) |
| --- | --- | --- | --- |
| Sex: f/m | N=24 (42.1%) / N=33 (57.9%) | N=46 (52.9%) / N=41 (47.1%) | p=0.206, Phi=-0.105 |
| Age-related motor experience (y) | M=36.74, BCa95% [33.72 39.75]<br>SD=11.91, BCa95% [10.06 13.36]<br>Min=8, Max=63 | M=10.10, BCa95% [9.49, 10.76]<br>SD=3.21, BCa95% [2.26 4.03]<br>Min=4, Max=27 | **Mdiff=26.63, p<0.001**<br>**BCa5% [23.40 29.78]**<br>**d=3.38 95%CI [2.86 3.89]** |
| Height (m) | M=1.74, BCa95% [1.71 1.77]<br>SD=0.10, BCa95% [0.08 0.11]<br>Min=1.42, Max=1.94 | M=1.42, BCa95% [1.39 1.45]<br>SD=0.13, BCa95% [0.11 0.14]<br>Min=1.12, Max=1.75 | **Mdiff=0.32, p<0.001**<br>**BCa95% [0.29 0.36]**<br>**d=2.73 95%CI [2.27 3.19]** |
| Weight (kg) | M=75.60, BCa95% [72.14 79.18]<br>SD=15.23, BCa95% [12.42 17.59]<br>Min=47, Max=123 | M=33.01, BCa95% [30.41 35.69]<br>SD=11.40, BCa95% [10.20 12.61]<br>Min=12, Max=64 | **Mdiff=42.59, p<0.001**<br>**BCa95% [38.02 47.23]**<br>**d=3.27 95%CI [2.76 3.77]** |
| BMI (kg/m2) | M=24.80, BCa95% [23.99 25.67]<br>SD=3.42, BCa95% [2.79 3.98]<br>Min=18.9, Max=35.5 | M=15.88, BCa95% [15.11 16.66]<br>SD=3.25, BCa95% [2.89 3.60]<br>Min=9.3, Max=24.2 | **Mdiff=8.92, p<0.001**<br>**BCa95% [7.86 9.99]**<br>**d=2.69 95%CI [2.23 3.14]** |
| Variability in balancing performance, EC (SD dCoP; mm/s2) | M=213.99, BCa95% [185.04 242.53]<br>SD=120.05, BCa95% [102.96 132.83]<br>Min=61.62, Max=487.91 | M=201.26, BCa95% [188.84 232.83]<br>SD=106.20, BCa95% [90.95 119.97]<br>Min=51.47 513.64 | Mdiff=3.73, p=0.84<br>BCa95% [-35.27 42.79]<br>d=0.03 95%CI [-0.30 0.37] |
| Pairing: Same sex / different sex | N=32 (56.1%) / 25 (43.9%) | N=44 (50.6%) / 43 (49.4%) | p=0.513, Phi=-0.055 |
| Difference in age-related motor experience (y) | M=15.75, BCa95% [12.23 19.04]<br>SD=15.55, BCa95% [13.89 16.86]<br>Min=-16, Max=40 | M=-9.83, BCa95% [-13.02 -7.01]<br>SD=14.15, BCa95% [12.55 15.40]<br>Min=-40, Max=6 | **Mdiff=25.58, p<0.001**<br>**BCa95% [20.92 30.52]**<br>**d=1.74 95%CI [1.35 2.13]** |
| Height difference (m) | M=0.15, BCa95% [0.10 0.19]<br>SD=0.19, BCa95% [0.16 0.22]<br>Min=-0.20, Max=0.67 | M=-0.10, BCa95% [-0.13 -0.06]<br>SD=0.19, BCa95% [0.164 0.22]<br>Min=-0.67, Max=0.33 | **Mdiff=0.24, p<0.001**<br>**BCa95% [0.19 0.31]**<br>**d=1.25 95%CI [0.89 1.62]** |
| Weight difference (kg) | M=21.49, BCa95% [14.27 28.29] | M=-14.08, BCa95% [-18.90 -9.19] | **Mdiff=35.57, p<0.001** |
|  | SD=28.80, BCa95 [24.09 33.32]<br>Min=-62, Max=78 | SD=23.63, BCa95% [19.79 26.82]<br>Min=-78, Max=29 | **BCa95% [27.46 43.92]**<br>**d=1.38 95%CI [1.01 1.75]** |
| BMI difference (kg/m2) | M=4.84, BCa95% [3.05 6.46]<br>SD=6.96, BCa9% [5.73 8.12]<br>Min=-16.20, Max=16.20 | M=-3.17, BCa95% [-4.3 -1.99]<br>SD=5.64, BCa95% [4.93 6.31]<br>Min=-16.00, Max=8.90 | **Mdiff=8.01, p<0.001**<br>**BCa95% [5.93 10.01]**<br>**d=1.29 95%CI [0.93 1.66]** |
| Difference in balancing skill, EC (mm/s2) | M=0.29, BCa95% [0.07 0.52]<br>SD=0.90, BCa9% [0.71 1.05]<br>Min=-0.78, Max=3.15 | M=0.24, BCa95% [0.07 0.45]<br>SD=0.88, BCa95% [0.70 1.05]<br>Min=-0.76, Max=3.60 | Mdiff=0.05, p=0.68<br>BCa95% [-33.60 49.92]<br>d=0.05 95%CI [-0.28 0.39] |
| Benefit of IPT, EC (relative change due to IPT, EC (mm/s2)) | M=-70.57, BCa95% [-102.24 -35.51]<br>SD=125.13, BCa95% [106.78 141.30]<br>Min=-339.27, Max=188.40 | M=-79.12, BCa95%[-100.62 -58.88]<br>SD=104.40, BCa95% [82.57 119.99]<br>Min=-382.64, Max=163.76 | Mdiff=8.54, p=0.66<br>BCa95% [-30.70 45.32]<br>d=0.08 95%CI [-0.26 0.41] |
| Benefit of IPT (percentage sway change due to IPT, EC (%)) | M=-15.40, BCa95% [-31.20 3.64]<br>SD=67.57, BCa95% [46.10 84.80]<br>Min=-85.27, Max=216.08 | M=-30.05, BCa95% [-38.75 -22.31]<br>SD=39.94, BCa95% [33.03 45.74]<br>Min=-84.83, Max=82.96 | Mdiff=14.65, p=0.17<br>BCa95% [-4.50 34.88]<br>d=0.28 95%CI [-0.06 0.61] |

Proportional difference in balancing skill: (balancing skill self - balancing skill other) / (balancing skill other);
Percentage benefit of IPT: (balance performance with IPT - balance performance wo IPT) / balance performance wo IPT)\*100

**Figure S1.**
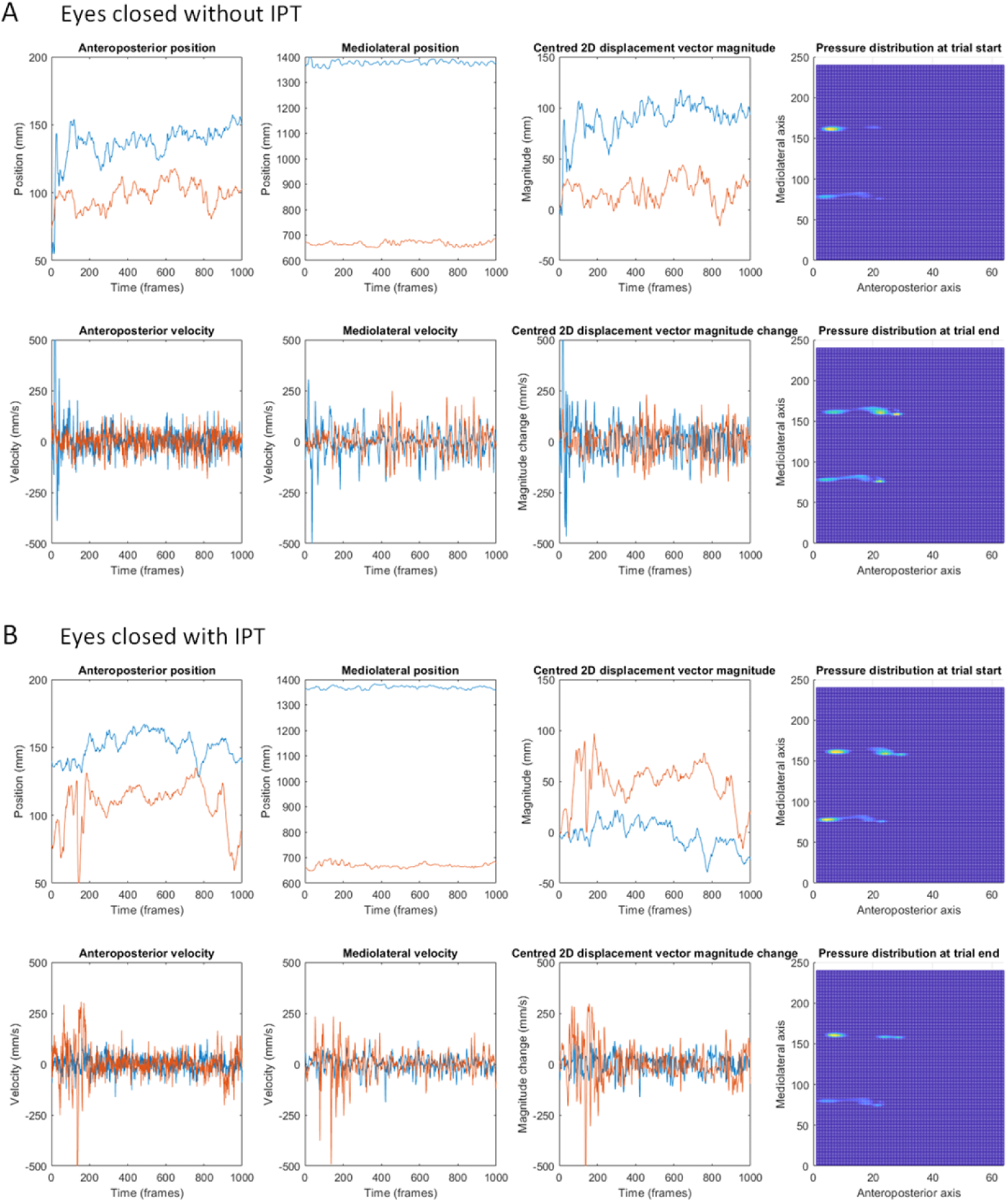
Illustrative data traces showing two individuals during side-by-side single-legged stance on a barometric platform with (A) Eyes closed without interpersonal touch and with (B) Eyes closed and simultaneous interpersonal touch. The participant on the left of the pair is shown as a line in Blue, the participant on the right as a line in Orange. The 2D displacement vector magnitude is the resultant of the anteroposterior and mediolateral positions. IPT: interpersonal touch.

**Figure S2.**
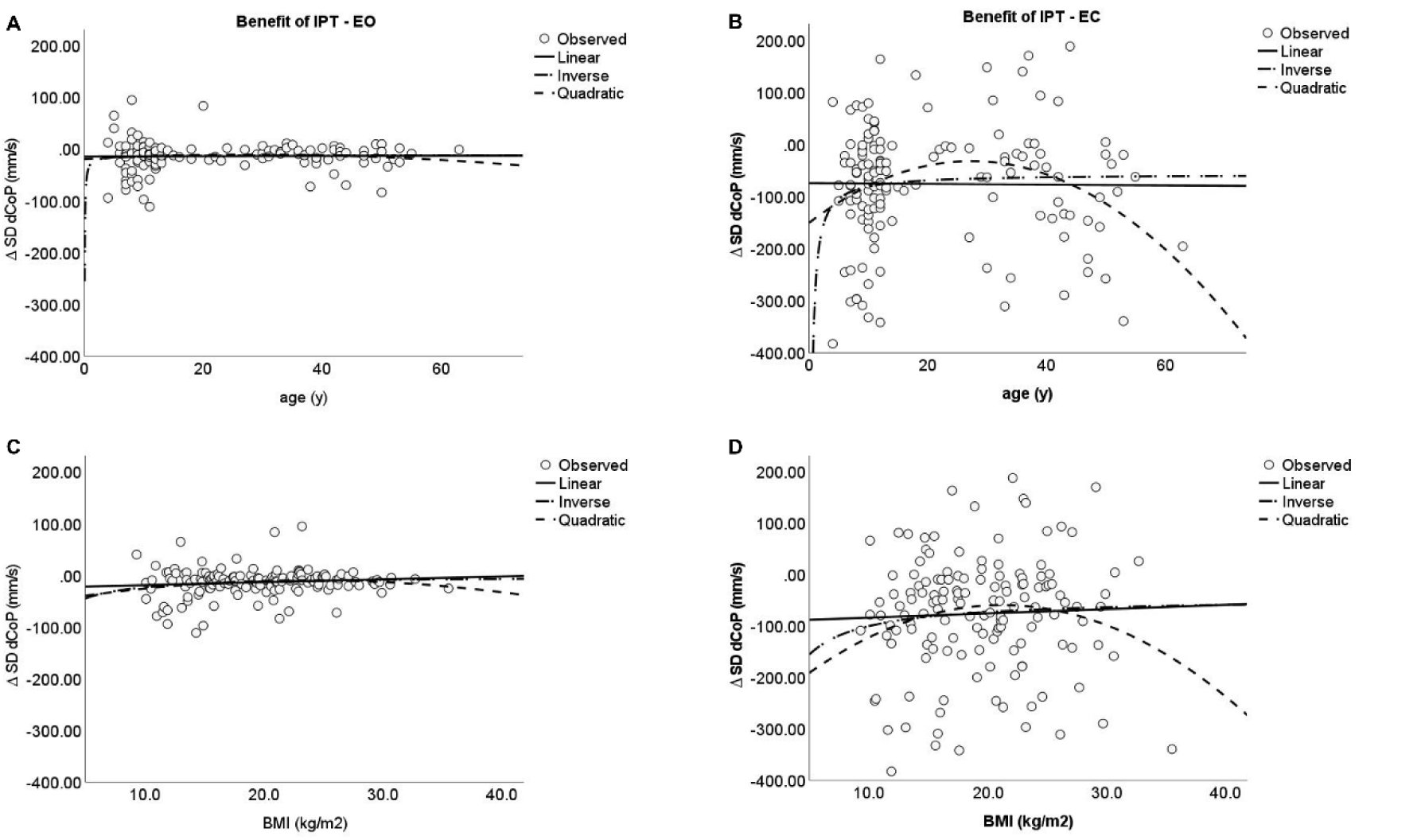
Curve fitting results for age (left) and BMI (right) with the benefit of IPT in Eyes open condition (EO; left) and Eyes closed condition (EC; right).

**Figure S3.**
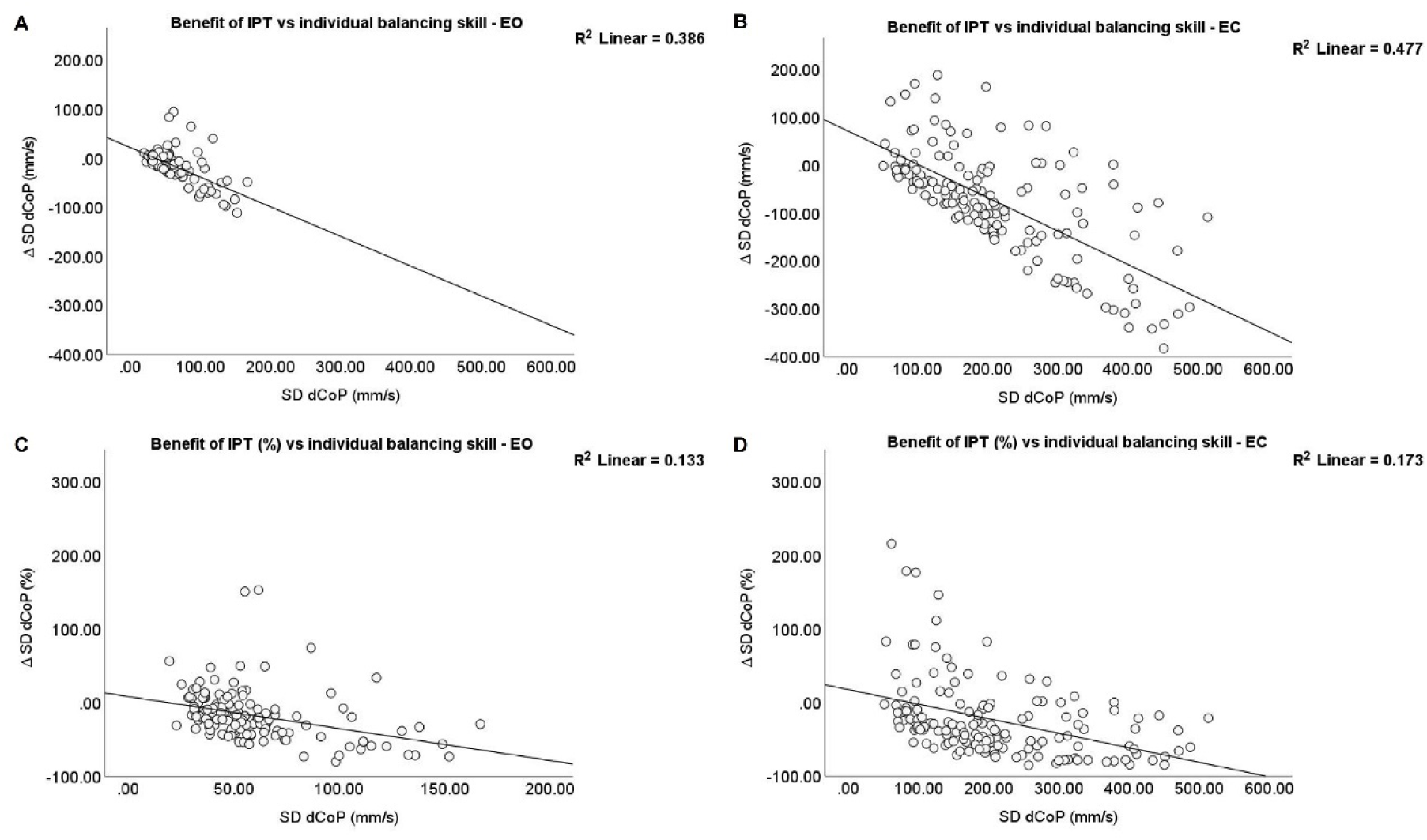
Scatterplot of the benefit of IPT (y-axis) in relation to an individual’s balancing skill (x-axis). A more negative delta SD dCoP (top) and percentage change in SD dCoP (bottom) indicate a greater sway reduction. EO: Eyes open, EC: Eyes closed; IPT: interpersonal touch. In the bootstrapped bivariate Pearson correlation analysis (N=1000, Seed=2021, BCa95%) a stronger correlation was observed for an individual’s balancing skill with the relative benefit of IPT (EO: r=-0.62, p<0.001, BCa95%CI [−0.78 −0.43]; EC: r=-0.69, p<0.001, BCa95% [−0.72 −0.61]) compared to with the percentage benefit of IPT (EO: r=-0.36, p<0.001, BCa95%CI [−0.52 −0.22], EC: r=-0.42, p<0.001, BCa95%CI [−0.50 −0.34]).

**Figure S4.**
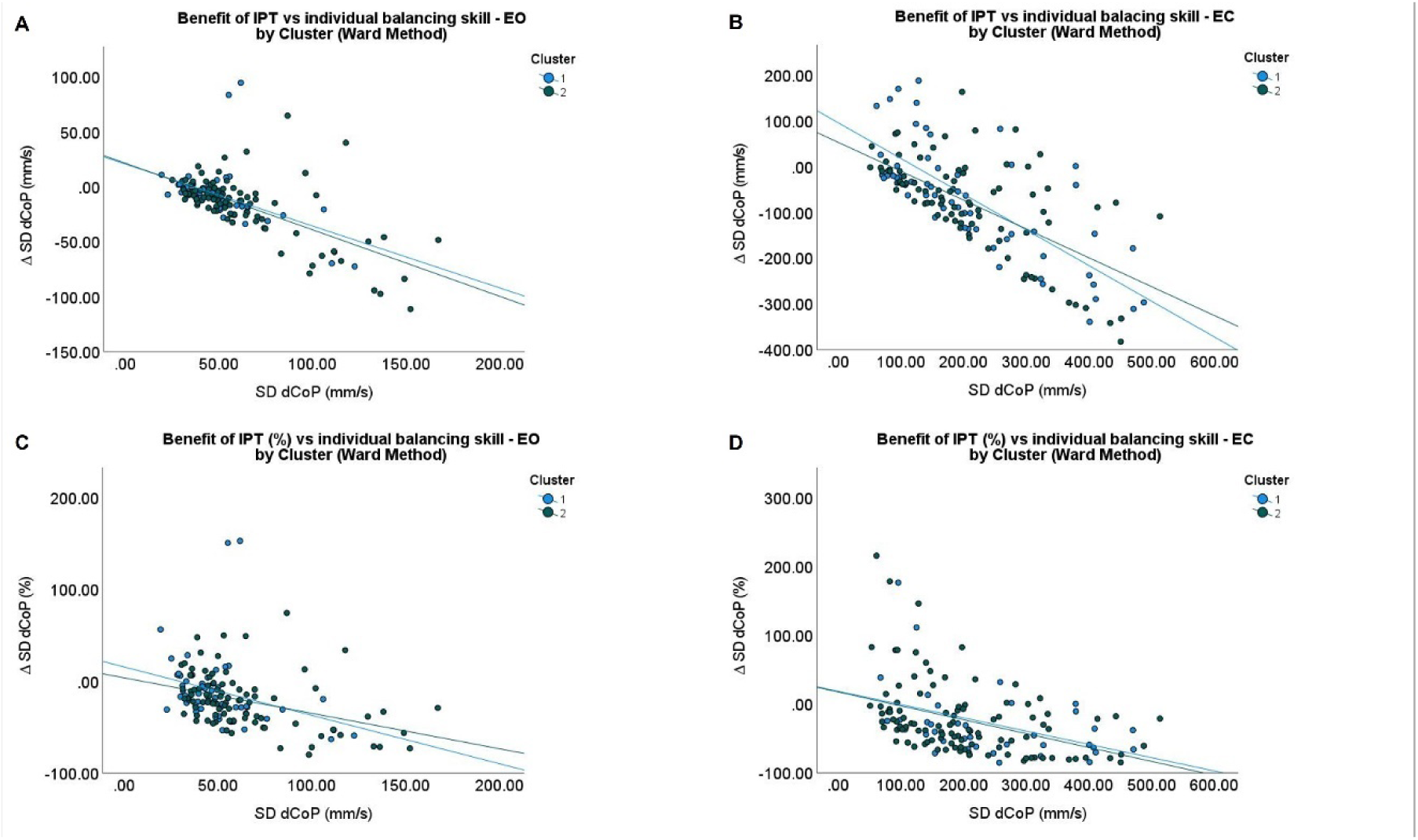
Scatterplot of an individual’s balancing skill and interindividual differences in balancing skills with the benefit of IPT dependent on the personalised performance cluster assignment.

The bootstrapped bivariate Pearson correlation analysis of the relative benefit of IPT across the whole group showed a moderate to high correlation with the interindividual differences in balancing skills (EO: r=-0.45, p<0.001, BCa95%CI [−0.58 −0.30]; EC: r=-0.54, p<0.001, BCa95%CI [−0.65 −0.40]). Bootstrapped bivariate Pearson correlations, did not reveal a significant relationship of an individual’s balancing skill (SD dCoP, no IPT) with age-related motor experience (r=-0.12, p=0.161, BCa95% [−0.28 0.05]), motor developmental potential (mean-centred age inverse) (r=-0.04, p=0.643, BCa95% [−0.11 0.03]) or extreme BMI (mean-centred BMI squared) (r=0.08, p=0.33, BCa95%CI [−0.07 0.27]) for Eyes open condition. Though, there was a significant low relationship of an individual’s balancing skill with height (r=-0.29, p<0.001, BCa95%CI [−0.45 −0.14]), weight (r=-0.25, p=0.002, BCa95%CI [−0.40 −12]) and BMI (r=-0.28, p<0.001, BCa95%CI [−0.42 −0.14]). Also, for the Eyes closed condition, age-related motor experience (r=0.02, p=0.847, BCa95%CI [−0.18 0.21]) and motor developmental potential (r=-0.06, p=0.505, BCa95%CI [−0.17 0.13]) were not significantly correlated with an individual’s balancing skill. In this condition only extreme BMI (r=0.18, p=0.031, BCa95%CI [− 0.01 0.35]) showed a tendency for a low correlation, not however, height (r=-0.11, p=0.194, BCa95%CI [−0.29 0.09]), weight (r=-0.04, p=0.641, BCa95%CI [−0.23 0.15]) and BMI (r=-0.06, p=0.504, BCa95%CI [−0.25 0.15]).

However, the curve estimation for Eyes open condition showed the inverse relationship (R2=0.09, p<0.001) to be the best fit for describing the relationship an individual’s balancing skill with age-related motor experience compared to a linear relationship (R2=0.01, p=0.16), followed by the quadratic relationship (R2=0.06, p=0.012). For the relationship of BMI with body sway, the quadratic relationship (R2=0.11, p<0.001) better represented the relationship than the linear estimation (R2=0.08, p<0.001), though the linear relationship was also significant. For the relationship of an individual’s balancing skill and age-related motor experience during EC condition, the inverse (R2=0.03, p=0.045) and quadratic relationships (R2=0.05, p=0.029) revealed to be significant and to explain a greater amount of variance than the linear relationship (R2=0.000, p=0.847). The relationship of an individual’s balancing skill with BMI in the Eyes closed condition showed only the quadratic relationship (R2=0.05, p=0.040) to be significant and explained a greater amount of variance than the linear relationship (R2=0.003, p=0.504). To be consistent across sensory conditions and due to the low number of individuals aged >50 years (N=9) and >55 years (N=2), the inverse relationship was further used in the analysis for both Eyes open and Eyes closed.

**Figure S5.**
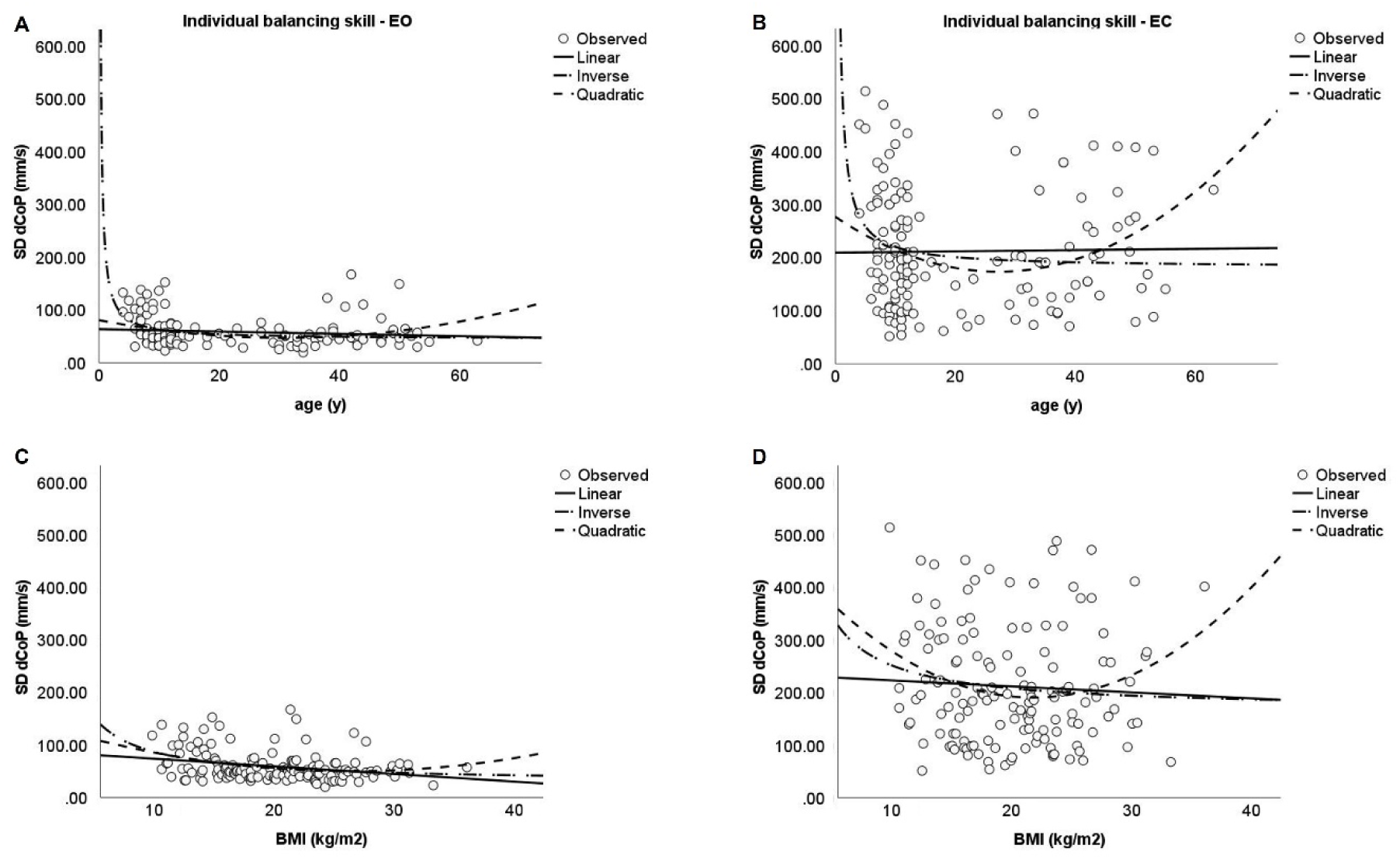
Curve fitting results of an individual’s balancing skill with Eyes open (left) and Eyes closed(right) based on age-related motor experience (top) and BMI (bottom).

In the serial mediation analysis for the Eyes open condition, 18% of the variance of an individual’s balancing skill (SD dCoP) was explained by height, BMI and the full serial mediation of the effect of age via height, weight, and BMI. Greater body sway was explained by a lower body height (B=-174.61, BootMean=-169.49, BootSE=77.86, Boot95%CI [−304.31 −0.42]), β =-1.21, and lower BMI (B=-6.45, BootMean=-6.40, BootSE=2.24, Boot95%CI [−11.03 −1.80], β =-1.24). Age (B=0.56, BootMean=0.59, BootSE=0.36, Boot95%CI [−0.06 1.43], β =0.30), weight (B=2.14, BootMean=2.08, BootSE=1.11, Boot95%CI [−0.26 4.08], β =1.84), sex (B=-0.75, BootMean=-4.23, BootSE=5.12, Boot95%CI [−15.09 5.42], β =-0.07), age mean-centred inverse (B=-0.75, BootMean=-2.00, BootSE=6.25, Boot95%CI [−19.83 8.07], β =-0.01) and BMI mean-centred squared (B=0.10, BootMean=-0.10, BootSE=0.12, Boot95%CI [−0.34 0.15], β =-0.13) did not contribute significantly to the model. The effect of age on body sway was mediated via height (B=-1.75, BootSE=0.78, Boot95%CI [−3.05 −0.00], β =-0.93, BootSE=0.44, Boot95%CI [−1.74 −0.00]), height and BMI (B=0.83, BootSE=0.30, Boot95%CI [0.23 1.44], β =0.44, BootSE=0.17, Boot95%CI [0.12 0.78]), weight and BMI (B=-0.85, BootSE=0.38, Boot95%CI [−3.12 −0.50], β =-0.45, BootSE=0.21, Boot95%CI [−0.95 −0.11]), and via height, weight and BMI (B=-0.99, BootSE=0.64, Boot95%CI [−1.67 −0.26], β =-0.99, BootSE=0.35, Boot95%CI [−1.67 −0.26]).

For the Eyes closed condition, 14% of the variance of body sway was explained by height, weight, BMI and sex, as well as by the full serial mediation of the effect of age on body sway via height, weight, and BMI. Geater body sway was explained by a lower height (B=-955.55, BootMean=-934.89, BootSE=306.57, Boot95%CI [−1538.72 −346.46], β =-1.70), a greater weight (B=13.59, BootMean=13.35, BootSE=4.56, Boot95%CI [4.57 21.95], β =3.00), a lower BMI (B=-30.06, BootMean=-29.29, BootSE=10.63, Boot95%CI [−48.22 −6.77], β =-1.48) and for females (B=43.49, BootMean=43.54, BootSE=18.20, Boot95%CI [5.80 76.21], β =0.20). Age (B=0.73, BootMean=0.60, BootSE=1.25, Boot95%CI [−2.13 2.76], β =0.10), age mean-centred inverse (B=-20.52, BootMean=-20.45, BootSE=43.02, Boot95%CI[−118.82 76.80], β =-0.06), and BMI mean-centred squared (B=-0.73, BootMean=-0.74, BootSE=0.50, Boot95%CI [−1.75 0.21], β =-0.24) did not contribute significantly to the model. However, the effect of age was mediated via height (B=-9.59, BootSE=3.21, Boot95%CI [− 15.61 −3.35], β =-1.31, BootSE=0.40, Boot95%CI [−2.04 −0.48]), weight (B=5.66, BootSE=2.31, Boot95%CI [1.53 10.63], β =0.78, BootSE=0.30, Boot95%CI [0.22 1.39]), via height and weight (B=12.30, BootSE=4.46, Boot95%CI [3.81 21.35]), via height and BMI (B=3.87, BootSE=1.45, Boot95% CI [0.83 6.67], β =0.53, BootSE=0.19, Boot95%CI [0.12, 0.89]), via weight and BMI (B=-3.97, BootSE=1.61, Boot95%CI [−7.24 −0.91], β =-0.54, BootSE=0.21, Boot95%CI [−0.95 −0.13]), and via height, weight and BMI (B=-8.61, Boot 95%CI [−14.97 −1.80], β =-1.18, BootSE=0.42, Boot95%CI [−1.98 −0.26]).

Moreover, the variance of interindividual differences in balancing skills during Eyes open condition was explained to 11% by height difference and a mediated effect of age difference via height difference. A more positive interindividual difference in balancing skill was explained by a lower height difference (B=-1.83, BootMean=-1.77, BootSE=0.61, Boot 95%CI [−2.91 −0.50], β =-0.58) and the mediated effect of age difference via height difference (B=-0.02, BootSE=0.01, Boot95%CI [−0.03 −0.01], β =-0.44, BootSE=0.17, Boot95%CI [−0.76 −0.10]). Age difference (B=0.01, BootMean=0.01, BootSE=0.01, Boot95%CI [−0.00 0.03], β =0.30), weight difference (B=0.02, BootMean=0.01, BootSE=0.01, Boot95%CI [−0.00 0.03], β =0.66), BMI difference (B=-0.06, BootMean=-0.06, BootSE=0.04, Boot95%CI [−0.13 0.02], β =-0.60) and sex difference (B=-0.03, BootMean=-0.03, BootSE=0.12, Boot 95%CI [−0.25 0.19], β =-0.02) did not have a significant direct effect.

In the Eyes closed condition, interindividual differences in balancing skills were explained to 10% by height difference (B=-2.60, BootMean=-2.53, BootSE=0.81, Boot95%CI [−4.28 −1.02], β =-0.66), weight difference (B=0.05, BootMean=0.05, BootSE=0.02, Boot95%CI [0.02 0.08], β =1.78), and BMI difference (B=-0.16, BootMean=-0.16, BootSE=0.05, Boot95%CI [−0.25 −0.05], β =-1.30). Moreover, differently to the Eyes open condition, during Eyes open the effect of age difference on interindividual differences in balancing skills was fully mediated via height-, weight- and BMI differences (B=0.04, Boot 95%CI [−0.07 −0.01], β =-0.87, BootSE=0.31, Boot95%CI [−1.39 −0.29]), in addition to via height difference only (B=-0.02, BootSE=0.01, Boot95%CI [−0.04 −0.01], β =-0.50, BootSE=0.16, Boot95%CI [−0.84 −0.20]), via weight difference (B=0.02, BootSE=0.01, Boot95%CI [0.01, 0.04], β =0.50, BootSE=0.18, Boot95%CI [0.15 0.87]), via height and weight differences (B=0.04, BootSE=0.02, Boot95%CI [0.02 0.08], β =0.94, BootSE=0.31, Boot95%CI [0.32, 1.55]), via height and BMI differences (B=-0.02, BootSE=0.01, Boot95%CI [0.01, 0.03], β =-0.35, BootSE=0.12, Boot95%CI [0.10 0.57]), and via weight and BMI differences (B=-0.02, BootSE=0.01, Boot95%CI [− 0.04, −0.01], β =-0.46, BootSE=0.16, Boot95%CI [−0.77 −0.14]). Age difference (B=0.00, BootMean=0.00, BootSE=0.01, Boot95%CI [−0.01 0.01], β =0.02), and sex difference (B=-0.01, BootMean=-0.01, BootSE=0.14, Boot95%CI [−0.29 0.29], β =-0.00) did again not show any significant direct effect.

**Figure S6.**
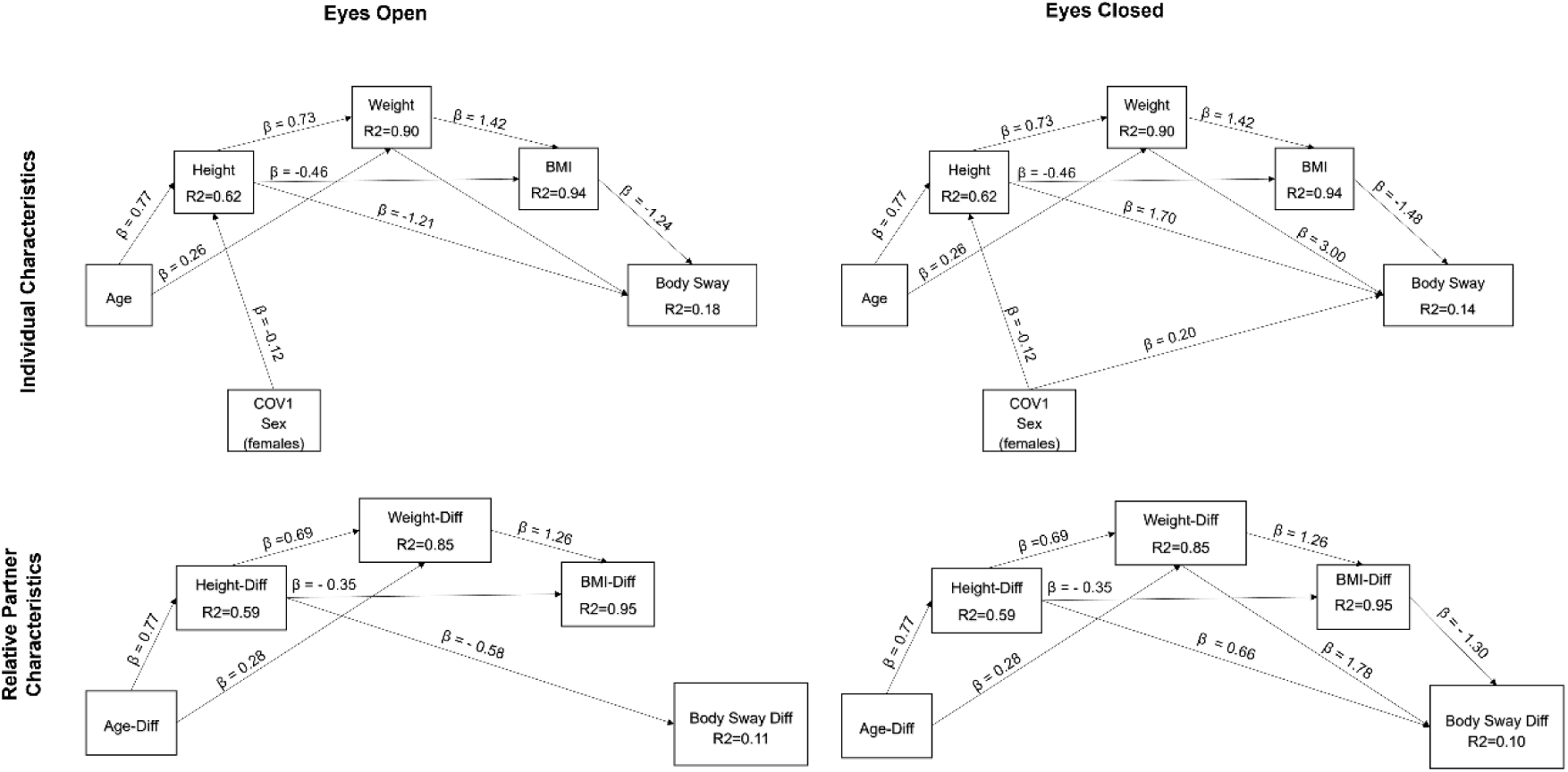
Statistical model of the influential factors on the benefit of IPT (relative difference in variability in balancing skills with IPT compared to without IPT) for Eyes open condition (top) and Eyes closed condition (bottom).

